# The Impact of Variant Calling on Substitution Mutational Signature Inference

**DOI:** 10.64898/2025.12.09.693327

**Authors:** Zichen Jiang, Jessica N. Au, Mariya Kazachkova, Marcos Díaz-Gay, Raviteja Vangara, Ludmil B. Alexandrov

**Affiliations:** Department of Cellular and Molecular Medicine, University of California San Diego, La Jolla, CA, USA; Department of Bioengineering, University of California San Diego, La Jolla, CA, USA; Moores Cancer Center, University of California San Diego, La Jolla, CA, USA; Bioinformatics and Systems Biology Graduate Program, University of California San Diego, La Jolla, CA, 92093, USA; Biomedical Sciences Graduate Program, University of California San Diego, La Jolla, CA, USA; Digital Genomics Group, Cancer Genomics Program, Spanish National Cancer Research Center, Madrid, Spain; Sanford Stem Cell Institute, University of California San Diego, La Jolla, CA, USA

**Author notes:** These authors contributed equally (Zichen Jiang, Jessica N. Au).

## Abstract

Identifying mutational signatures is a key component of cancer genomics studies, yet the influence of variant calling strategies on signature extraction has not been systematically evaluated. Here, we analyzed over 8,900 whole exomes from The Cancer Genome Atlas (TCGA) and over 1,800 whole genomes from the Pan-Cancer Analysis of Whole Genomes (PCAWG) consortium to assess how mutation callers shape *de novo* single-base substitution (SBS) signatures. We found that consensus calling yielded stable *de novo* signatures across reference genomes and pipeline versions, whereas individual callers introduced false-positive SBSs that manifested as artifactual signatures that were reproducibly detected by three independent signatures extraction tools. A minimal consensus approach requiring agreement between only two variant calling algorithms effectively removed these artifacts while preserving true biological signal. Together, these results establish consensus variant calling as essential for robust inference of *de novo* SBS mutational signatures and provide practical guidelines for distinguishing genuine mutational processes from technical artifacts.

## INTRODUCTION

Somatic mutations are present in all healthy and cancer cells^1,2^. They arise from the activities of endogenous and exogenous mutational processes, many of which leave characteristic patterns of mutations, termed *mutational signatures*^3,4^. Over the past decade, mutational signature analysis has become an integral part of genomics studies^5^, as it provides a systematic framework to identify and understand the processes that generate somatic mutations. Biologically, these signatures capture the footprints of diverse mechanisms, including DNA replication errors^4^, defective DNA repair pathways^4,6^, enzymatic activities such as APOBEC cytidine deaminases^6^, and exposures to environmental mutagens such as ultraviolet light^7^, tobacco smoke^8^, or chemotherapy^9^. By linking patterns of mutations to their underlying causes, mutational signature analysis not only deepens our understanding of genome maintenance and carcinogenesis but also provides insights into normal cellular aging^1^ and potential therapeutic vulnerabilities^10^.

From a technical perspective, detecting mutational signatures in cancer samples generally follows a predefined sequence of steps. First, cancers and their matched normal tissues (most often blood) are subjected to next-generation sequencing^11^. The sequencing reads are aligned to a reference genome and somatic mutations are then identified by comparing the tumor to its matched normal with one or more bioinformatics tools, thereby filtering out shared germline variants^11^. The resulting catalog of somatic mutations is converted into a mutational matrix^12^ that summarizes the distribution of mutation types across all examined samples. Computational tools implementing factorization methods, such as nonnegative matrix factorization (NMF)^13–15^, are then used to decompose this matrix into putative mutational signatures. Finally, these extracted signatures are compared against reference catalogs, such as the Catalogue of Somatic Mutations in Cancer (COSMIC) mutational signatures database^16^, to identify known signatures and detect any potentially novel ones.

Current mutational signature analyses typically treat all cancer genome mutations as equivalent, failing to account for potential biases introduced by reference genomes and variant calling algorithms. As an example, The Cancer Genome Atlas (TCGA) relied on the MC3 project^17^ to use whole-exome sequencing (WES) data aligned to GRCh37 to detect consensus somatic mutations in a “2+/3” manner, stipulating a mutation should be identified by at least two out of three callers (MuTect^18^, MuSE^19^, VarScan2^17,20^). When the Genomic Data Commons (GDC) subsequently reanalyzed the same sequencing data^21^ using an updated bioinformatics pipeline—which included alignment to GRCh38, updated annotations, and consensus 2+/3 variant calling with MuTect2^22^ as well as newer sub-versions of MuSE and VarScan2—the finalized set of single base substitutions (SBSs) differed by 21%^21^. Similar variability is also evident in cancer genomics studies that have utilized whole-genome sequencing (WGS) data. The Pan-Cancer Analysis of Whole Genomes (PCAWG) consortium used a consensus 2+/4 approach requiring support from at least two of four callers (MuTect, MuSE, DKFZ^23,24^, CaVEMan^25^), whereas subsequent reanalysis of the same samples by the Hartwig Medical Foundation with its SAGE^26^ variant caller produced different mutation counts and profiles. While the choice of reference genome and mutation-calling algorithm impacted which mutations are detected^21,27,28^, especially those with low variant allele frequency (VAF)^29^, their effect on the extraction and interpretation of mutational signatures remain unexplored.

In this study, we systematically evaluated how the choice of reference genome and variant calling approach influenced the analysis of mutational signatures in WES and WGS data. We focused on single base substitution (SBS) signatures, the most prevalent and widely used class of mutational signatures. Leveraging WES data from >8,900 TCGA tumors common to both MC3 legacy and GDC harmonized public releases (**Fig. S1**) and WGS data from >1,800 PCAWG tumors processed by five variant callers (**Fig. S2**), we assessed the robustness of SBS mutational signature extraction and quantified the impact of technical factors on the biological interpretation of mutational processes.

## RESULTS

### Mutational signatures and mutational profiles

A set of SBSs detected in a cancer genome can be summarized into a mutational profile, which classifies mutations according to substitution type and sequence context^4^. For simplicity, substitutions are usually represented using the pyrimidine base of the Watson-Crick base pair (e.g., C>A instead of C:G>T:A)^4^. The simplest scheme, SBS-6, groups mutations into six categories (C>A, C>G, C>T, T>A, T>C, and T>G). Greater resolution is achieved with SBS-96, which incorporates the immediate 5’ and 3’ flanking bases, yielding 96 trinucleotide-based classes and forming the most widely used representation in mutational signature analyses. Both SBS-288 and SBS-1536 build upon SBS-96 by adding additional layers of context information. SBS-288 separates mutations in intergenic versus genic regions, and within genic regions further distinguishes between the transcribed and un-transcribed strands. This helps to capture transcriptional strand asymmetries introduced by processes such as transcription-coupled repair^5,12,14^. SBS-1536 instead expands to a pentanucleotide context (two bases on each side), generating 1,536 classes that can help disentangle signatures with otherwise similar SBS-96 patterns^12,14^. A generalized scheme that combines the features of SBS-288 and SBS-1536 is SBS-4608, which incorporates both transcriptional strand bias and pentanucleotide context to provide the highest-resolution mutational patterns.

*De novo* extraction of mutational signatures typically analyzes hundreds of tumor samples^4,14,15^, each summarized into a mutational profile defined under a chosen classification schema. These schemas range in resolution from SBS-6 (generally too coarse for practical use)^4,12^ to SBS-96 (currently the most widely used schema^14^). Some studies^30,31^ have also employed SBS-288 or SBS-1536 to capture additional biological context, but because the COSMIC reference catalog^16^ is defined in SBS-96, results ultimately need to be converted back to SBS-96 space for comparison and consistency across studies. In practice, similarity between mutational profiles or extracted signatures is almost always quantified using cosine similarity^13^: values above 0.95 are generally considered near-perfect matches, values between 0.85 and 0.95 indicate a good match, while values below 0.85 suggest non-matching profiles. For context, two random nonnegative vectors have an expected average cosine similarity of 0.75^32^.

Computational tools for *de novo* mutational signature extraction have previously sought to account for the impact of mutation calling variability on the extracted signatures. For example, SigProfilerExtractor applies Poisson resampling prior to NMF to mitigate unsystematic errors and improve the robustness of the resulting solutions^15^. This tool also automatically determines the optimal number of operative mutational signatures (denoted by *k*) in a dataset, reconstructs their patterns, and assigns contributions of each signature to individual samples. Importantly, SigProfilerExtractor has been extensively benchmarked and consistently shown to outperform other *de novo* extraction approaches^15^. However, these strategies can only buffer against random noise and cannot correct for systematic biases, such as artifactual mutations introduced by specific variant calling algorithms, which may propagate into mutational profiles and distort signature extraction. Moreover, to date, no systematic comparison has evaluated the effectiveness of signature extraction across different variant callers or genome builds, leaving an open question about the generalizability of its performance under distinct variant calling pipelines.

### Consensus WES mutations yield consistent SBS profiles

While mutational signature extraction depends on mutation classification schemes and computational safeguards against error, the accuracy of the underlying mutation calls remains paramount. To evaluate how technical choices shaped SBS analyses, we compared 8,908 TCGA tumors shared between the legacy MC3 and harmonized GDC releases (**Figure S1**), which differed in both reference genome and consensus 2+/3 variant calling pipeline. This design enabled a direct assessment of how such differences affected downstream SBS mutational signature analyses. Despite the differences in processing the underlying WES data, over 51% of the tumors exhibited virtually identical SBS-96 sample profiles (cosine similarity ≥0.95; **Figures 1A and 1B**). Importantly, this concordance increased with tumor mutation burden, with tumors harboring more mutations exhibiting stronger similarity between pipelines (Spearman’s *ρ*=0.833; p<0.0001; **Figure 1B**).

**Figure 1.**
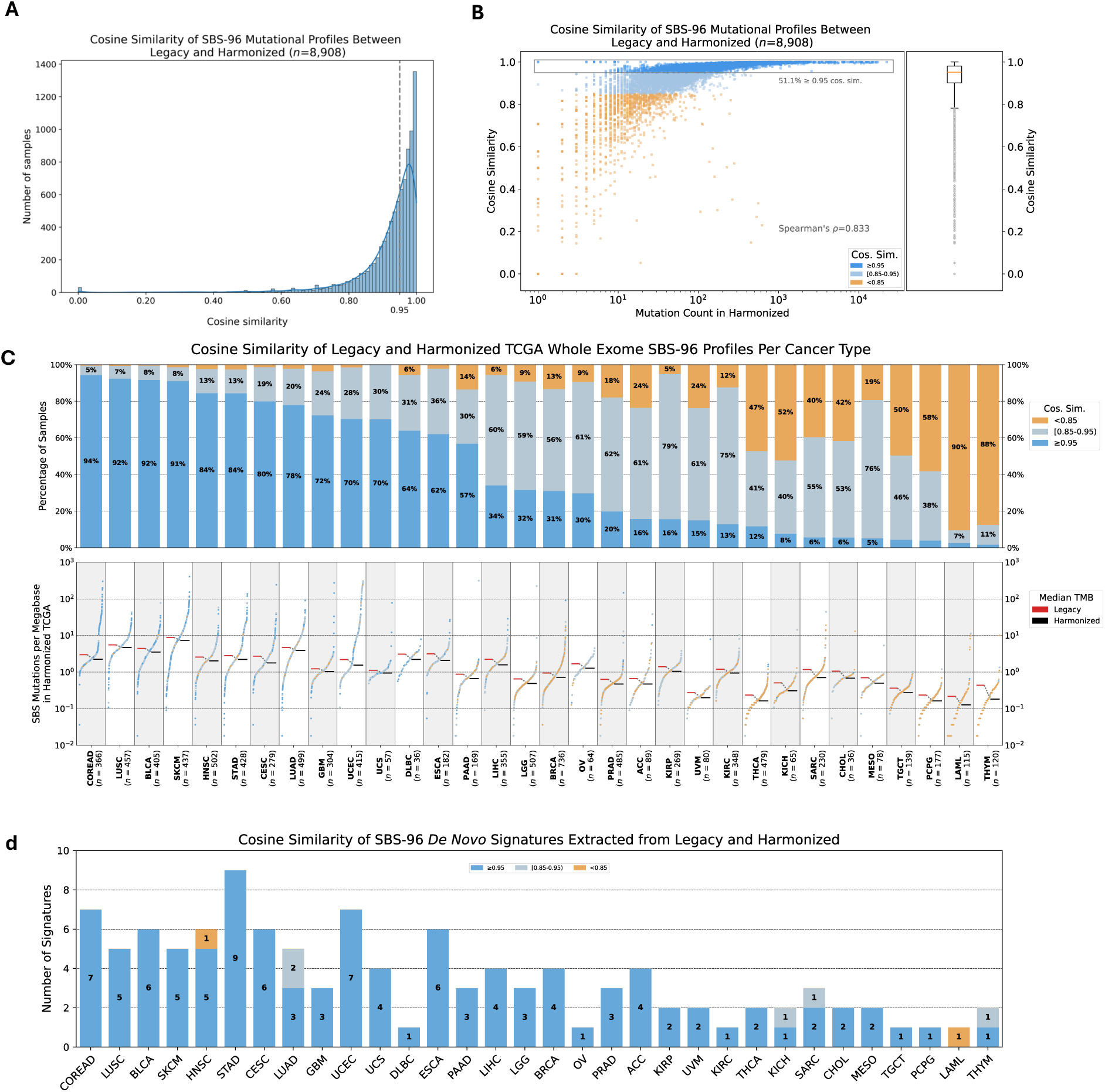
Impact of TCGA harmonization of consensus variant calls on the downstream whole exome SBS-96 mutational signature extractions. **(A)** Histogram displaying the number of TCGA primary tumors at each bin for SBS-96 sample profile cosine similarity between legacy and harmonized consensus calling datasets. A kernel density estimate curve is overlaid on the histogram. The gray vertical dashed line marks the cosine similarity value of 0.95, denoting near-perfect profile concordance. **(B)** *Left*: Scatter plot relating the sample SBS mutation count (x-axis; log-scale) to profile concordance (y-axis) between legacy and harmonized TCGA datasets. Each dot corresponds to an individual tumor. Samples with cosine similarity below 0.85 are highlighted in orange. *Right*: Box plot summarizing the overall distribution of cosine similarities. **(C)** *Top*: Stacked bar plot showing the proportion of samples per cancer type within three tiers of cosine similarity (≥0.95, [0.85,0.95), and <0.85). Cancer types are sorted by the proportion of samples in the ≥0.95 bin. *Bottom*: Snake plot showing the mutational burden (SBSs per 55 Megabases (Mb); assuming that a whole-exome sequence allows calling mutations in approximately 55 MB) for each sample in the harmonized dataset, grouped by cancer type. Horizontal lines indicate the median mutational burden for the harmonized (black) and legacy (red) datasets. (D) Stacked bar plot showing the number of nearly identical, similar, and dissimilar *de novo* extracted SBS-96 signatures between legacy and harmonized datasets for each cancer type, at a forced number of signatures, which is often the suggested *k*. The default normalization method of Gaussian mixture model was used. Dissimilar signatures that were altered by the harmonization process are highlighted in orange. i.e., Neck Squamous Cell Carcinoma (HNSC) and Acute Myeloid Leukemia (LAML).

To evaluate the reliability of mutational profiles, we applied Poisson resampling to the harmonized GDC SBS-96 sample profiles (**Figure S3; Methods**). This procedure would simulate stochastic sampling noise and account for unsystematic errors in a profile, illustrating that the confidence in a pattern is strongly dependent on mutation burden. Across 1,000 Poisson resamplings for each of the 8,908 TCGA tumors, we found that profiles defined by only a few mutations were inherently unstable, and resampling made this uncertainty explicit by producing profiles that deviated substantially from the observed pattern. By contrast, profiles supported by hundreds or thousands of mutations remained highly robust, with resampled versions nearly identical to the original. This analysis revealed that WES samples required at least 383 SBSs to maintain a stable mutational profile under random error. Above this threshold, 95% of resampled profiles achieved a cosine similarity ≥0.95 between the observed and resampled profiles.

### Consensus WES mutations yield consistent SBS signatures

While a slight majority of tumors (51.1%; **Figures 1B and 1C**) exhibited almost identical SBS-96 mutational profiles between the legacy and the harmonized releases, a substantial subset of the exomes exhibited striking discrepancies. We asked whether the discordant samples would drive divergence of *de novo* mutational signatures extracted from the two consensus-based datasets. Remarkably, despite the variability at the sample-level, the cohort-level extracted signatures between the two TCGA releases were largely consistent. First, across cancer types, SigProfilerExtractor suggested the same number of operative signatures (*k*) in 24 of 32 TCGA cancer types for both the GDC and MC3 releases (**Figure S4**). In seven of the remaining eight cancer types, the inferred *k* differed by only one between the two datasets, while for uterine corpus endometrial carcinoma (TCGA-UCEC) it differed by two. Second, when the number of signatures was fixed to the one detected in the GDC release, the individual signatures extracted from both datasets were highly similar to each other, with cosine similarities consistently ≥0.95 (**Figure 1D**). For example, in TCGA’s breast invasive carcinoma (TCGA-BRCA) cohort, consensus calling pipelines from the two releases yielded four stable signatures with near-perfect concordance. The notable exception was acute myeloid leukemia (TCGA-LAML), where low sample counts and mutation burdens resulted in both dissimilar sample profiles and mutational signatures. Overall, both the GDC and MC3 releases employed a consensus variant calling approach, defining somatic mutations only when detected by two or more callers. Using this strategy, mutational profiles and especially mutational signatures were remarkably stable despite differences in genome builds, consensus pipelines, and the overall number of somatic mutations.

### Individual mutation callers yield variable signatures in WES data

Having established the robustness of mutational signatures under consensus variant calling across reference genomes and consensus pipelines, we next investigated the effect of individual variant callers in WES cancers. We focused on the harmonized TCGA-BRCA WES data and compared SBSs identified by a “2+/3” consensus strategy (*i.e.*, mutations identified by at least two of the three bioinformatics tools) with those derived from the individual callers: MuTect2, MuSE, and VarScan2. Each individual caller’s distinct algorithm produced discordant mutation sets. For instance, MuTect2 employs a Bayesian model for local haplotype assembly^22^, which confers high sensitivity for detecting low variant allele frequency (VAF) mutations (**Figure 2A**). Consequently, this led to the identification of over 47,814 MuTect-exclusive SBSs not captured by the other two callers (**Figure 2B**). The choice of individual variant calling algorithm substantially affected the SBS-96 profile of a given tumor, with no pair of tools yielding more than 20% of samples with cosine similarity ≥0.95 (**Figure 2C**). This sample-level discordance in TCGA-BRCA WES profiles resulted in divergent *de novo* SBS-96 signatures derived from VarScan2 and MuSE mutation sets, but surprisingly not from MuTect2. Compared to the consensus 2+/3 baseline *de novo* signatures, both VarScan2 and MuSE yielded a distinct signature that could represent a potentially either novel process or technical artifacts **(Figure 2D)**.

**Figure 2.**
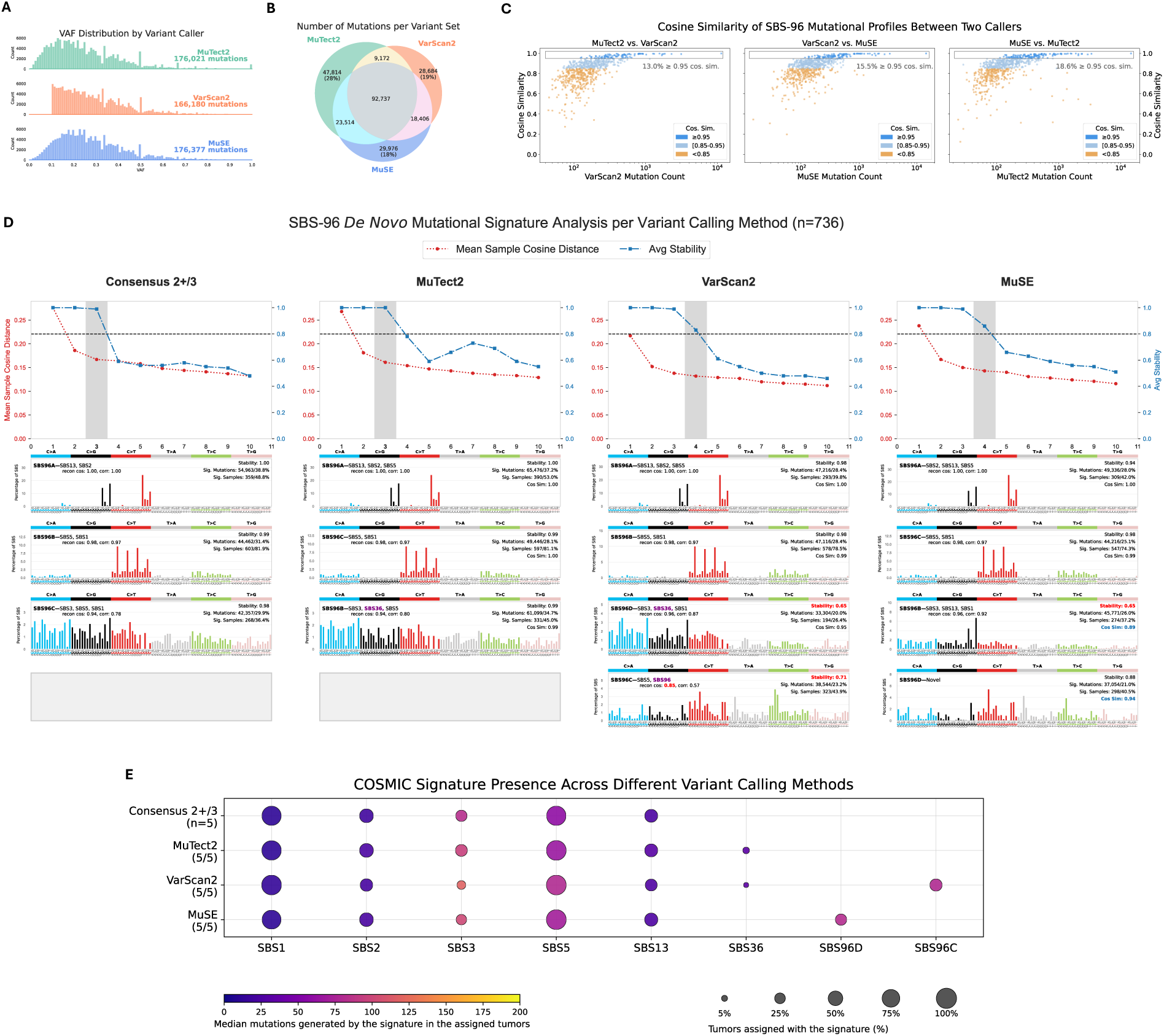
Influence of variant callers on harmonized TCGA-BRCA whole-exome SBS-96 mutational signatures. **(A)** Histogram plot of variant allele frequencies (VAFs) for all SBSs identified by three individual algorithms used in the consensus calling of harmonized TCGA-BRCA samples. A hard VAF cutoff of 10% is applied by VarScan2 algorithm’s “PASS” filter. **(B)** Venn-diagram of mutations showing the number of exclusive SBSs identified by each mutation calling algorithm. **(C)** Pairwise comparison of SBS-96 mutational profiles generated from different callers. Each scatter plot relates the sample SBS mutation count (x-axis; log-scale) to the cosine similarity (y-axis) between two variant calling algorithms. Each dot corresponds to an individual tumor. Samples with cosine similarity below 0.85 are highlighted in orange. **(D)** *Top*: Signature selection plots for SBS-96 *de novo* extraction, showing the average stability of each solution with a given number of signatures. Suggested number of solutions is highlighted by a grey vertical bar. Samples in the input matrix with over 400 SBSs were normalized before NMF factorization. *Bottom*: The signatures of the suggested solution for each caller. Gray rectangles denote that no similar signature can be found (cosine similarity <0.85). On the top right corner of each signature plot, low stability scores are highlighted in red and the number of assigned samples and mutations are indicated. On the top left, poor decomposition is highlighted in red (reconstruction cosine similarity <0.90). *D*e *novo* signatures that cannot be decomposed at all are denoted as “novel”. Reference signatures in the decomposition result that are not typically seen in breast cancers are labeled in purple. **(E)** Dot plot showing the decomposition of *de novo* signatures from each mutation caller into COSMICv3.4 reference signatures and their exposure in the variant call sets. Number of active COSMICv3.4 signatures is noted in parenthesis, with the consensus 2+/3 extraction result used as benchmark. When reference decomposition is poor (cosine <0.90) or cannot be decomposed at all, the *de novo* signature itself is used to quantify the exposure.

We then mapped the *de novo* extracted signatures from each variant caller to a modified COSMICv3.4 reference set that excluded ultraviolet-related signatures, polymerase-epsilon associated hypermutator signatures, and other signatures not relevant to breast tissue (**Methods**). In TCGA-BRCA, signatures extracted from individual callers yielded mutational patterns absent from the COSMIC reference, including SBS96C from VarScan2 (323/736 samples; 23.2% of total mutations) and a highly comparable signature, SBS96D, from MuSE (298/736 samples; 40.5% of mutations; **Figure 2D and 2E**). Both signatures likely reflected mutation calling artifacts, as the mutations exclusive to each caller occurred in regions masked by RepeatMasker^33^ and Tandem Repeats Finder^34^, had sequencing strand bias, and/or had insufficient coverage (<30 reads) (**Figure S5**). Particularly, masked regions are difficult to map and prone to variant calling errors, making putative somatic events identified by a single caller hard to distinguish from technical artifacts^35^. Indeed, signatures extracted solely from caller-exclusive mutations closely matched the artifactual signatures identified in each caller’s full mutation set. These TCGA exome results showed individual variant detection tools can produce spurious artifactual mutational signatures due to caller-exclusive false positives, emphasizing the value of consensus variant calling.

### Individual mutation callers yield variable signatures in WGS data

We next evaluated the impact of variant calling strategies in WGS data by examining 1,857 PCAWG samples^28^ that were processed by the original set of four variant callers (i.e., MuTect, MuSE, DKFZ, and CaVEMan) as well as SAGE^26^ (**Figure S2**). Specifically, 2,376 of the 2,700 samples in the original PCAWG cohort were included in the SAGE re-analysis by Martínez-Jiménez et al.^26^. From there, we retained only cancer types with at least 50 samples to ensure adequate statistical power for signature analysis, resulting in a final set of 1,857 samples for analysis (**Methods**). Over 88% of samples from individual callers had cosine similarity over 0.95 when compared to the PCAWG consensus calls (2+/4 of the original set of four variant callers), but some samples with high mutation burden still exhibited different profiles (**Figure 3A**). As in TCGA WES, the most variable cancer types were those with low median mutational burdens, defined as fewer than one SBS per megabase, including medulloblastoma and pilocytic astrocytoma (**Figure 3B**). However, unlike WES, *de novo* signature extraction from WGS data produced variable results across all cancer types, regardless of their median mutational burden (**Figure 3C**). Furthermore, enforcing a consistent number of signatures was challenging, as each caller’s suggested solution deviated substantially from the consensus (**Figure S6**). These results indicated that the algorithmic differences among variant callers could produce distinct sets of somatic mutations and sample profiles in WGS data, which in turn might yield caller-specific signatures or tissue-irrelevant patterns.

**Figure 3.**
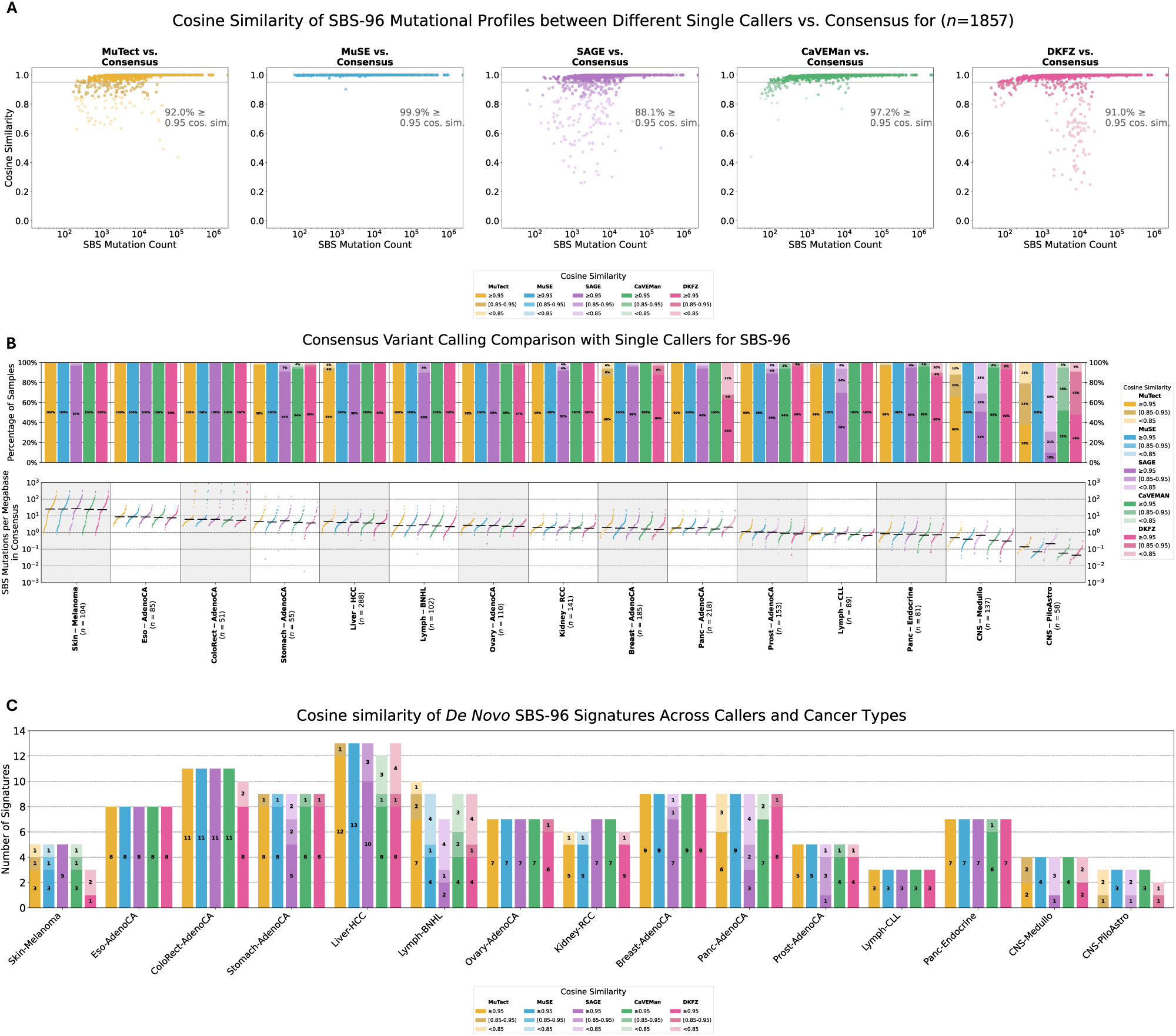
Comparison of individual versus PCAWG consensus variant calling for whole genome mutational signature extractions. **(A)** Scatter plots showing the cosine similarity of SBS-96 sample profiles between PCAWG consensus (2+/4) and five variant callers, in relation to SBS mutation count. Each scatter plot compares a sample’s SBS mutation count (x-axis; log-scale) and the cosine similarity (y-axis) between a mutation calling algorithm and the consensus mutational calling approach. Each dot corresponds to an individual tumor. Samples with cosine similarity below 0.85 are highlighted in orange. **(B)** *Top*: Stacked bar plot showing the proportion of samples per cancer type per variant caller within three tiers of cosine similarity (≥0.95, [0.85,0.95), and <0.85). Cancer types are sorted by the proportion of samples in the ≥0.95 bin averaged across the five variant callers. *Bottom*: Snake plot showing the mutational burden (SBSs per 2,800 Megabases (Mb); assuming that a whole-genome sequence allows calling mutations in approximately 2,800 MB) of each sample determined by each individual variant caller. Black horizontal lines indicate the median mutational burden for the datasets. **(C)** Stacked bar plot showing the number of *de novo* SBS-96 signatures extracted from individual variant callers that are nearly identical, similar, or dissimilar relative to consensus (2+/4) solution (see online methods for solution picking in WGS extractions). The number of caller-specific signatures in a cancer type is highlighted in lighter hues.

### Single variant callers produce artifactual mutational signatures in WGS data

To assess the biological impact of single-caller variability, we analyzed 185 PCAWG breast adenocarcinoma (Breast-AdenoCA) whole genomes using mutation calls from five independent tools: MuTect, MuSE, SAGE, CaVEMan, and DKFZ. The number of somatic SBSs identified varied across algorithms, ranging from 1,277,780 by DKFZ to 1,532,806 by MuTect (**Figure 4A**), with only 959,172 mutations shared by all five tools (**Figure 4B**). Additionally, MuSE and MuTect had the largest number of exclusive mutations—those not called by any other algorithm—at approximately 200,000 each (**Figure 4B**). Despite this discordance in individual mutation calls, pairwise comparisons of aggregated SBS-96 profiles involving these callers showed that over 80% of samples displayed nearly identical patterns between them (cosine similarity ≥0.95; **Figure 4C**).

**Figure 4.**
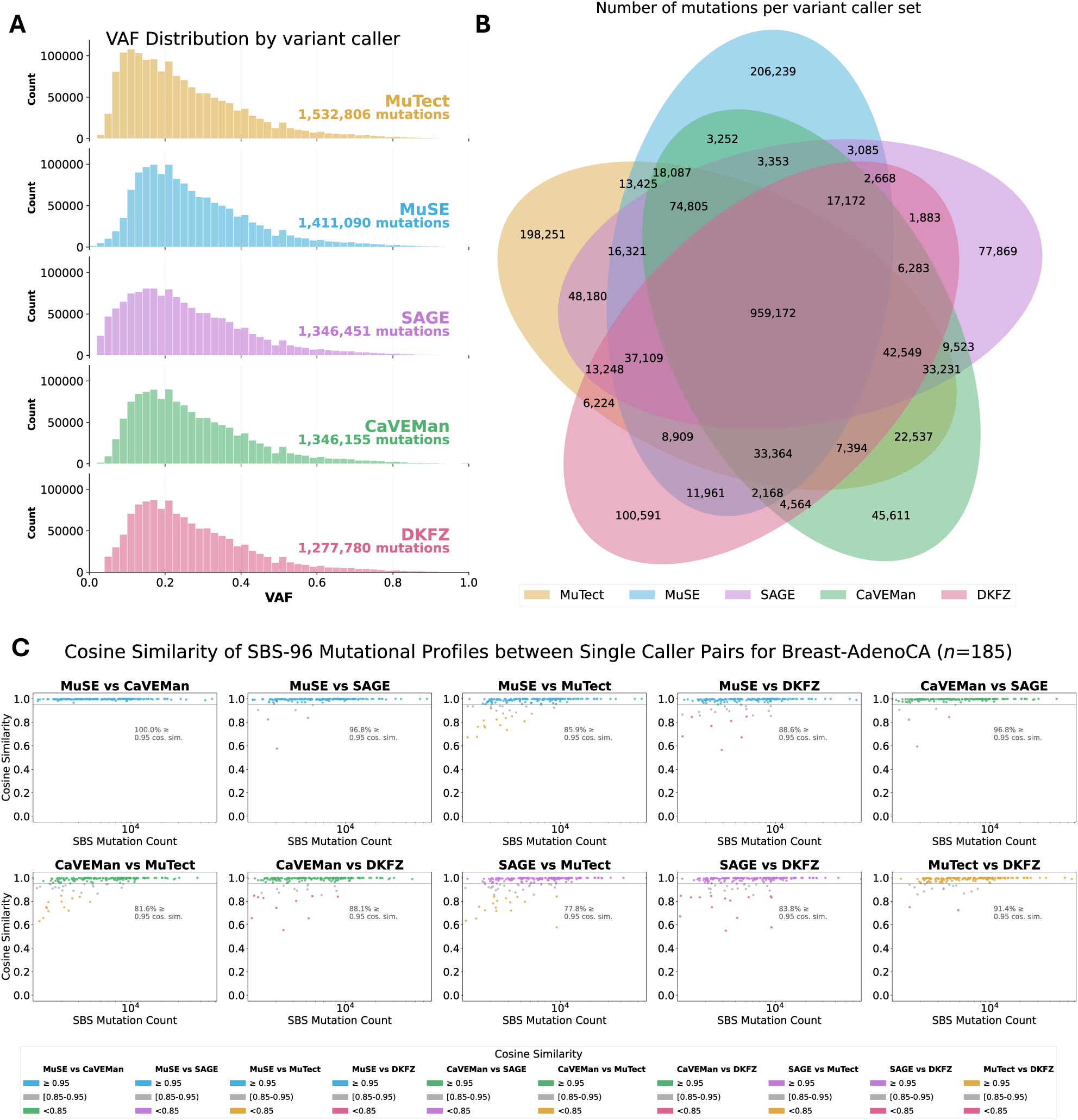
SBS-96 sample profiles in 185 WGS Breast AdenoCA using different variant callers. **(A)** Histogram plot of variant allele frequencies (VAFs) for all SBSs identified by MuTect (yellow), MuSE (blue), SAGE (purple), CaVEMan (green), and DKFZ (pink) in 185 PCAWG Breast-AdenoCA whole genomes. **(B)** Venn diagram quantifying both the unique and the intersections of SBS mutations called by the five distinct callers. **(C)** Pairwise scatter plots relating cosine similarity of SBS-96 sample profiles generated from two callers (y-axis) to SBS mutation count (x-axis; log-scale). Points are colored to highlight discordance. Samples with high similarity between two callers (cosine ≥0.95) match the color of the first caller listed in the plot title, while dissimilar samples (cosine <0.85) match the second. Samples with intermediate concordance (cosine 0.85-0.95) are colored gray. A gray rectangle outline in each plot marks the nearly identical samples.

However, this apparent concordance at the profile level masked critical differences when signatures were extracted *de novo*. Extraction from the consensus 2+/5 set of somatic mutations (comprising mutations identified by at least two of the five independent callers) served as our baseline and yielded nine *de novo* signatures (**Figure 5A**). These included signatures associated with known biological processes: clock-like aging (SBS1/5)^36^, APOBEC activity (SBS2/13)^6^, homologous-recombination deficiency (HRD; SBS3/8)^4^, and reactive oxygen species (ROS; SBS18)^37^. It also identified signatures of unknown etiologies: SBS17a/b, SBS28, SBS34, SBS37, SBS39, and SBS40a^38^ (**Figure 5B**).

**Figure 5.**
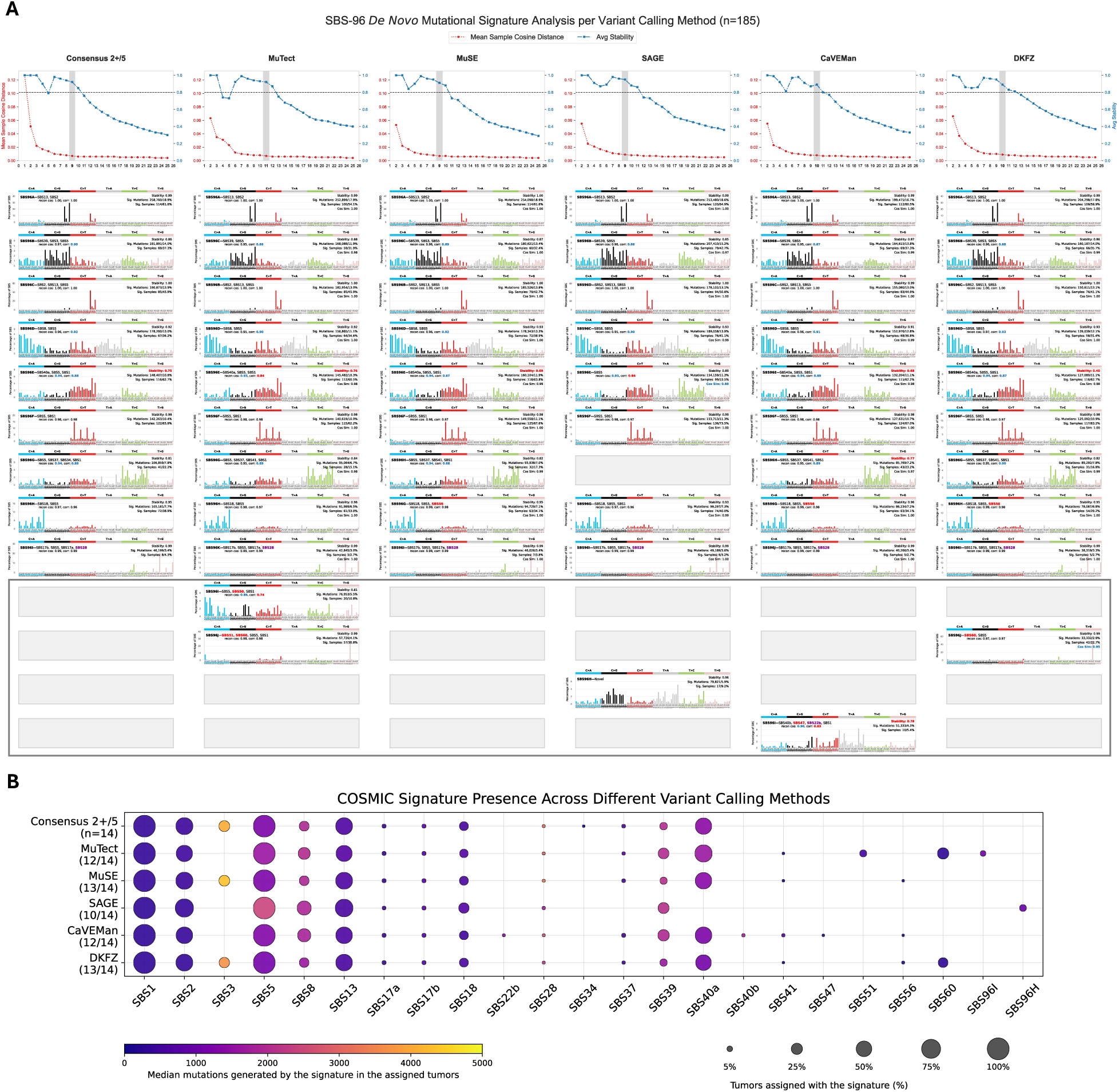
SBS-96 mutational profile differences between variant caller specific mutation sets in PCAWG Breast-AdenoCA whole genomes. **(A)** *Top*: Signature selection plots from *de novo* extraction using consensus mutations identified by custom consensus strategy (2+/5 callers) and mutations called by five individual callers. Suggested number of solutions is highlighted by a grey vertical bar. *Bottom:* Suggested *de novo* signatures extracted from each variant calling method. Gray rectangles signify that no similar signature was found (cosine similarity below 0.85). The region marked by a grey line encompasses artifactual signatures identified a specific variant caller. For each signature plot, low stability scores and poor decompositions (reconstruction cosine similarity <0.90) are highlighted in red text. COSMICv3.4 reference signatures are atypical for breast cancer in the decomposition result are labeled in purple. *De novo* signatures that could not be decomposed are denoted as “novel”. The number of assigned samples and mutations is indicated for each signature. **(B)** Dot plot showing the decomposition of *de novo* signatures from each caller into COSMICv3.4 reference signatures and their exposure in the variant call sets. Number of reference signatures is noted in parenthesis with the consensus 2+/5 result used as benchmark. When reference decomposition was poor (cosine below 0.90) or could not be decomposed, the *de novo* signature itself is used to quantify the exposure.

Compared to the consensus 2+/5 baseline, individual variant callers introduced false positives. MuTect, for example, reproduced nine of the consensus signatures but extracted two additional stable signatures not found in the baseline (SBS96I and SBS96J). The evidence for these two MuTect-specific signatures was clear. SBS96I generated above 5% of mutations called by MuTect and was active in over 10% of the 185 samples. SBS96J accounted for 4% of the mutations in the cohort and was found in over 30% of the samples. These *de novo* signatures were artifacts. SBS96I imperfectly decomposed to a sequencing artifact COSMICv3.4 SBS50 (reconstruction cosine similarity 0.86), and SBS96J neatly decomposed into two other sequencing artifacts, SBS51 and SBS60 (reconstruction cosine similarity 0.98)^14^. MuTect’s SBS96J was recapitulated in the DKFZ extraction (cosine similarity 0.95).

Similarly, SAGE produced a unique signature (SBS96H) not found by any other callers, accounting for >81,000 mutations (6% of SAGE calls) across 17 samples (9.2%; **Figure 5A**). Further investigation revealed this signature was artifactual. SAGE called a disproportionately higher number of low VAF mutations compared to other callers (**Figure 4A**), with these mutations comprising up to 50% of total variants in some samples exhibiting SBS96H. Applying a simple VAF filter (≥0.05) and re-performing *de novo* signature extraction completely eliminated SAGE SBS96H from the suggested solution and recovered all nine signatures that are nearly identical to the consensus 2+/5 baseline, implying that SBS96H arose from potentially low-confidence calls rather than genuine biological signal.

To find the possible source of these variant caller-specific signatures, we manually inspected the alignment files from samples whose caller-specific profiles showed high cosine similarity to the MuTect- and SAGE-specific signatures (MuTect’s SBS96I, MuTect’s SBS96J, and SAGE’s SBS96H). First, we focused on the MuTect-exclusive mutations in the following four prominent contexts of MuTect’s SBS96I: G[C>G]A, G[C>G]C, G[C>T]T, and T[T>C]C. Mutations in these contexts consistently exhibited read bias, with alternate reads mapping exclusively to one strand of the reference genome (**Figure S7A**). Similarly, read bias artifacts drove MuTect to call mutations in the G[T>G]G context, producing MuTect’s SBS96J that resemble a mixture of SBS51 and SBS60 (**Figure S7B**). For SAGE’s SBS96H, inspection of variants within the characteristic peaks (e.g., C[C>G]T) revealed that the vast majority were supported by only a single and completely overlapping pair of reads that span one short DNA insert of ∼100 bp (**Figure S7C**). This is consistent with sequencing noise rather than genuine somatic variants. We found that removing SAGE mutations supported by only two alternate reads eliminated the SAGE SBS96H artifact and recovered all nine signatures indistinguishable from the consensus 2+/5 baseline. This removal strategy proved to be as effective as ≥0.05 VAF filtering. To further confirm their technical origin, extractions restricted to caller-exclusive mutations recapitulated the three artifactual MuTect and SAGE signatures originally identified in the full call sets. These findings demonstrate that a single caller’s algorithmic biases can systematically misidentify sequencing noise as genuine mutations, and caller-exclusive mutations contribute to stable yet spurious mutational signatures.

### Consensus variant calling eliminates technical artifacts while preserving biological signal in WGS

Given that both MuTect and SAGE produced distinct artifactual mutational signatures, we investigated whether a minimal consensus approach could systematically eliminate these technical signatures while preserving genuine biological signal without requiring additional filtering. We found that intersecting any two variant callers effectively mitigated the caller-specific artifacts that confounded mutational signature analysis in the SBS-96 context (**Figure S8A**). We established a ground-truth set of signatures derived from the mutations present in all five callers. Our consensus strategies (2+/3 and 2+/5 callers) and the PCAWG method (2+/4 callers) accurately recapitulated these ground-truth signatures. Remarkably, while signatures derived independently from MuTect or DKFZ call sets produced an artifactual SBS60-like signature observed in Figure 5, this artifact was eliminated when extracting from the intersection of their mutation calls (minimum cosine similarity with the ground truth: 0.997; **Figure S8A**). Moreover, a similar disappearance of artifacts was observed in the *de novo* signatures extracted from the MuTect and SAGE intersect, demonstrating that a minimal consensus of just two callers was sufficient to eliminate strong, caller-specific artifacts. Of note, while MuSE-derived signatures exhibited high concordance with the ground truth (cosine ∼0.997), this was driven by MuSE’s reduced sensitivity to mutations below 0.10 VAF as illustrated in Figure 4a. This aligns with the inherent characteristic of our intersection-based ground truth, which also filters out low-VAF variants exclusive to a caller, rather than reflecting superior artifact suppression of MuSE. Furthermore, intersecting MuSE with just one additional caller (e.g., MuTect) yielded signatures indistinguishable from the five-caller consensus signatures, increasing the minimum cosine similarity from 0.975 to 0.993, highlighting the benefit of a two-caller consensus strategy.

### *De novo* mutational signature extraction of WGS using MuSiCal and SignatureToolsLib

To rule out whether these artifactual signatures are specific to a single signature extraction algorithm, we performed the same analysis using two other *de novo* extraction tools: MuSiCal^39^ and SignatureToolsLib^40^. The *de novo* signatures extracted from the 2+/5 consensus mutation calls were highly similar across all three methods (cosine similarity >0.85) (**Figure S9**), demonstrating that different extraction algorithms converged on comparable mutational signatures.

Most importantly, both MuSiCal and SignatureToolsLib independently detected the same caller-specific artifactual signatures identified by SigProfilerExtractor. The SAGE-specific signature (SBS96H) and MuTect-related artifacts (SBS96I and SBS96J) were consistently extracted across all three tools with highly similar profiles (cosine similarity >0.95; **Figure S10 and S11**). The robust detection of these artifacts across independent extraction tools, despite differences in their underlying mathematical frameworks, strongly indicated that these signatures originated from the variant callers themselves rather than from algorithmic artifacts introduced during signature extraction.

### Variability in signatures from extended SBS contexts

Finally, we assessed whether these findings were consistent when using higher-resolution SBS contexts. The change in genome build and consensus-calling pipeline overhaul largely did not have an impact on TCGA WES SBS-192 *de novo* signature analysis (**Figure S12)**. Analyzing the PCAWG WGS data with the SBS-288 (**Figure S13**), SBS-1536 (**Figure S14**), and SBS-4608 (**Figure S15**) extended contexts revealed that the divergence between single callers became even more pronounced. As the number of mutational channels increased, the median cosine similarity of sample profiles between individual callers decreased, indicating greater discordance at the sample level (**Figure S13A, S14A, and 15A**). We observed similar trends when re-analyzing PCAWG Breast-AdenoCA using MuSiCal as the SBS-288 signature extraction tool, indicating this phenomenon was not specific to SigProfilerExtractor. This suggested that when performing mutational signature analyses, the problem of single-caller variability persists in higher-definition analytical frameworks. However, we demonstrated that the use of any two variant callers resolved the issue of extracting artifactual *de novo* signatures in higher-resolution contexts such as SBS-288 (**Figure S16**).

## DISCUSSION

*De novo* mutational signature extraction has transformed our understanding of DNA damage and repair processes in cancer, yet the influence of upstream variant calling strategies on signature inference has remained largely unexamined. Across more than 8,900 TCGA whole exomes and 1,800 PCAWG whole genomes, we showed that technical choices made before signature extraction—particularly somatic variant calling—could shape downstream biological interpretation.

Our analyses demonstrated that consensus-based variant calling produced highly stable SBS signatures, even when reference genomes and caller versions differed substantially. In contrast, individual variant callers could introduce reproducible, algorithm-specific biases that manifested as stable artifactual signatures. These artifacts were reproducibly detected across at least three independent signature extraction frameworks (SigProfilerExtractor, MuSiCal, and SignatureToolsLib). They were also replicated by extracting from mutations exclusive to a specific caller. Manual inspection of sequencing alignment files showed that the mutations contributing to these signatures were poorly supported by sequencing reads and likely artifactual, confirming that these signatures arose from variant-caller-specific behavior rather than from the extraction algorithms themselves. Importantly, a minimal consensus strategy requiring agreement between any two callers effectively eliminated mutation caller-specific artifacts, even in mutational contexts beyond 96 channels. As a result, consensus variant calling should be considered a fundamental prerequisite for robust inference of SBS mutational signatures. For studies where consensus calling is impractical, *de novo* signatures extracted from single-caller analyses cannot be reliably distinguished from caller-specific artifacts without independent validation through orthogonal approaches. In the absence of such validation, stringent filtering such as excluding repetitive regions, strand-biased sites, weak read support, and low-quality base or mapping contexts, provides a partial safeguard. Particular caution is warranted when interpreting potentially novel SBS-96 signatures, which should be systematically checked against COSMIC references to exclude technical origins. These considerations become even more critical at higher-resolution SBS contexts (*e.g*., SBS-288, SBS-1536, SBS-4608), where caller discordance increases with the number of mutational channels and might amplify downstream variability.

Beyond pipeline-specific artifacts, our analyses highlighted that mutation burden itself is a determinant of signature reliability. Poisson resampling across 8,908 TCGA tumors showed that WES profiles supported by fewer than ∼383 SBSs was statistically unstable and diverged widely under resampling, whereas profiles exceeding this threshold remained robust. Ensuring an adequate mutation burden of 383 mutations should therefore be a standard criterion for WES signature extraction, alongside consensus calling, for reliable mutational signature analyses.

In conclusion, our results showed that upstream mutation calling exerted a strong and previously underappreciated influence on mutational signature inference. Consensus approaches and adequate mutation burden were critical for suppressing technical artifacts and ensuring that downstream analyses faithfully reflected underlying biological processes. In practice, this means that mutational signature studies should, whenever possible, rely on agreement between at least two independent variant callers. When consensus calling is not feasible, stringent filtering can provide a partial and alternative safeguard. Together, these practices help ensure that extracted mutational signatures represent true biological processes rather than artifacts generated by the algorithms used to detect mutations.

## METHODS

### Data Cohorts and Processing

#### The Cancer Genome Atlas (TCGA) WES Cohort

Somatic mutation data from whole-exome sequencing (WES) of primary tumors from The Cancer Genome Atlas (TCGA) were obtained from two distinct, large-scale processing efforts.

#### Legacy (MC3) Data

The public legacy Mutation Annotation Format (MAF) file (mc3.v0.2.8.PUBLIC.maf.gz) was downloaded from cBioPortal. This dataset was generated by the Multi-Center Mutation Calling in Multiple Cancers (MC3) project.^17^ The MC3 pipeline aligned BAM files to the human reference genome GRCh37 (hg19) and applied a consensus of three somatic variant callers to identify SBS, namely MuTect, VarScan2, and MuSE. A "majority rules" approach was used to generate a consensus 2+/3 call set, which was then subjected to a series of automated filters to remove common artifacts. For this study, only primary tumors and variants with a PASS filter status in the public MAF were utilized.

### Harmonized (GDC) Data

The harmonized TCGA MAF files were obtained from the NCI’s Genomic Data Commons (GDC) Data Portal, via the ISB Cancer Gateway in the Cloud (ISB-CGC) (release 39).^41^ This reprocessing effort represents a comprehensive overhaul of the original TCGA data analysis.^21^ All sequencing data were realigned to the human reference genome GRCh38 (hg38), and SBS somatic variants were called MuTect2, VarScan2, and MuSE. The outputs were then aggregated and filtered to produce a final consensus 2+/3 MAF file. Only primary tumors and variants with a PASS filter status in the public and controlled-access MAF were used in our analysis.^41^

### TCGA Sample Selection and Controlled-Access Data

For the direct comparison of consensus pipelines, we focused on a set of 8,908 primary tumor samples that had at least one PASS SBS in both the public MC3 and GDC call sets. A substantial proportion of the PASS mutations identified by an individual algorithm is not available to the public, so to fully investigate the impact of individual callers, we obtained controlled-access VCF files for 736 TCGA breast cancer (BRCA) primary tumors from the GDC Data Portal. For each sample, we created separate mutation sets based on all PASS variants identified by MuTect2, VarScan2, and MuSE individually. The TCGA-BRCA consensus 2+/3 is then custom made using their PASS variants.

### PCAWG Consensus Data

Controlled-access somatic VCF files were obtained from the International Cancer Genome Consortium (ICGC) Data Portal^42^ for 2,703 samples^42^. These variants were generated as part of the core Pan-Cancer Analysis of Whole Genomes (PCAWG) analysis, which aligned data to the GRCh37 reference genome.^28^

The consensus somatic variant calls were produced by a "2+/4" method, which required a variant to be confidently called by at least two of four different pipelines: the Broad Institute pipeline using MuTect v1.14, MuSE v1.0rc, the Sanger Institute pipeline using CaVEMan v1.5.1, and the German Cancer Research Center (DKFZ) pipeline using their in-house SBS calling workflow v0.1.19. VCFs were filtered for PASS mutations. We also analyzed variants labeled as "low support," which were called by only a single pipeline, to represent individual caller sets.

### Hartwig Medical Foundation (HMF) Recalled Data

A subset of the PCAWG WGS samples (*n* = 2,376) was independently re-analyzed by Martínez-Jiménez et al. using the Hartwig Medical Foundation’s in-house somatic variant calling pipeline, SAGE^26^. We obtained these recalled VCFs to enable a direct comparison between the SAGE and PCAWG consensus approach and its individual member algorithms on the same samples. Only PASS mutations were utilized.

### PCAWG Sample Selection

For the PCAWG WGS analysis, we focused on cancer types with more than 50 samples for which calls were available from the PCAWG consensus and all individual callers being compared. This resulted in a final cohort of 1,857 unique primary tumor samples across 15 cancer subtypes.

### Consensus calls for PCAWG WGS Breast-AdenoCA samples

Custom made consensus VCFs were generated from PASS mutations intersected from the full set of “low support” PCAWG data consisting of four callers: MuTect, MuSE, CaVEMan, and DKFZ, along with the PASS mutations from the recalled SAGE data. This was only performed for the 185 Breast-AdenoCA samples.

### Somatic Variant Calling Algorithms

The somatic variant calls used in this study were generated by several distinct algorithms and pipelines, each with a unique underlying methodology. A summary is provided in **Table S1**.

### Mutational Matrix Generation

All mutational matrices for signature analysis were generated using SigProfilerMatrixGenerator v1.3.3.^12^ For each analysis cohort (e.g., harmonized TCGA with consensus mutation calling, PCAWG Breast-AdenoCA with SAGE mutation calling), the corresponding set of VCF files was used as input, specifying the appropriate reference genome (GRCh37 for MC3/PCAWG; GRCh38 for GDC Harmonized) and set down-sampling to exome regions as false. We generated matrices for a comprehensive set of SBS mutational contexts: 96 (standard trinucleotide context), 192 (trinucleotides in transcribed regions that incorporate transcriptional strands), 288 (192 plus intergenic regions), 1536 (pentanucleotide context), and 4608 (1536 plus transcriptional strands and intergenic regions).

#### *De Novo* Mutational Signature Extraction

##### SigProfilerExtractor

*De novo* extraction of mutational signatures was performed using SigProfilerExtractor v1.2.1.^15^ The tool operates by decomposing the input mutational matrix (*M*) into two smaller matrices, representing the signatures (*S*) and their activities (*A*), such that *M ≈ S x A*. This is achieved through an NMFk-based approach that minimizes the Kullback-Leibler divergence^43^. Before NMF, the mutational catalog is Poisson resampled and subsequently then normalized to avoid extracting signatures that reflect random noise or the specific mutational profiles from a handful of very few hypermutated samples.^43^ The default option uses a Gaussian mixture model (GMM) to find a SBS mutation count that defines the hypermutating group, with the minimum being 9,600 SBSs. After we performed Poisson resampling simulation using the harmonized TCGA consensus data (**Figure S3**), we selected 400 SBSs as the normalization threshold going forward (e.g., for the TCGA-BRCA consensus 2+/3 versus individual caller analysis). For whole genomes, the vast majority of PCAWG cancer types exceed the minimum 9,600 SBSs required by GMM. Other SBS thresholds based on log_2_ transformation of total mutation counts or based on rescaling up to a maximum of 13,000 or and 30,000 SBSs did not affect the extraction results, so the default GMM was used as the normalization method for whole-genome analyses.

We ran SigProfilerExtractor to search for solutions with the number of signatures (*k*) ranging from 1 to 10 for WES or 1 to 25 for WGS. We increased the independent NMF replicates from the default 100 to 150 for WES and 250 for WGS. Reference and opportunity genomes are set corresponding to the input mutation processing pipeline. All other parameters are left as the default, such as random initialization of the S and A matrices.

The optimal number of stable signatures (*k*) from WGS was automatically determined by SigProfilerExtractor’s model selection procedure that uses a modified NMFk to be robust against noise^43^ and then employs an iterative process to balance low reconstruction error and low number of *de novo* signatures, while stipulating that the solution’s average silhouette score >=0.80 and the minimum stability >=0.20. WES solution selection does not include this iterative process.

#### SignatureToolsLib

The same set of 185 PCAWG Breast-AdenoCA samples’ SBS-96 mutational catalogue was analyzed with SignatureToolsLib (v2.4.6).^40^ We extracted mutational signatures for *k* between 2 and 20. As previously described^15^, we specified nrepeats=200, nboots=20, clusteringMethod=“MC”, filterBest_RTOL=0.001, filterBest_nmaxtokeep=10, and left all other parameters as the tool’s default. Importantly, SignatureToolsLib does not provide a suggested solution, so we chose the same *k* picked by SigProfilerExtractor, or a lower *k* that have average stability ≥0.80. Mutational contexts higher than SBS-96 are not supported by SignatureToolsLib.

#### MuSiCal

We analyzed the same Breast-AdenoCA samples using MuSiCal to extract SBS-96 and SBS-288 *de novo* signatures (v1.0.0).^39^ Following the approach outlined in the MuSiCal manuscript^39^, we used these default parameters: bootstrap=True, init=“random”, method=“mvnmf”, min_iter=1000, max_iter=100000, conv_test_freq=100, and tol=1e-8. We used all MuSiCal default parameters: n_replicates=20, normalize_X=True, bootstrap=True, init=“random”, method=“mvnmf”, min_iter=1000, max_iter=100000, conv_test_freq=100, and tol=1e-8. MuSiCal performs its own solution picking, so the number suggested n_components was utilized in our analysis.

### Manual Solution Picking

In TCGA WES consensus calls, 24 cancer types had the same number of signatures *k* in the automatically selected solutions from the legacy MC3 and harmonized GDC versions while seven other cancer types differed by one and uterine corpus endometrial carcinoma (TCGA-UCEC) differed by two (**Figure S4**). The number of true positive signatures can often be one or more smaller than the number *k* in the suggested solution, according to prior simulations^15^. To account for this variability, we manually chose the same *k* for the WES extractions from the two releases, which is immediately adjacent to their respective suggested *k* values.

In PCAWG consensus 2+/4 calls and individual callers, the number of extracted signatures shown is the minimum of the suggested solutions for consensus 2+/4 and each variant caller for each cancer type (**Figure 3C**). Using Skin-Melanoma as an example, the suggested solution reported by SigProfilerExtractor was *k* = 5 for the consensus 2+/4 mutations. We compared the cosine similarity of these five *de novo* signatures suggested solution as reported for each individual caller. The suggested solution for the MuTect-exclusive signatures was *k =* 7, then the stacked bar plot shows the top five of the seven signatures that are most similar to the consensus. The suggested solution for DKFZ-exclusive mutations was *k* = 3, so the most similar three signatures against the consensus *k* = 5 are shown.

### Signature Decomposition to COSMIC Reference Catalogue

To interpret the biological meaning of the de novo extracted signatures, we decomposed them into the reference signatures from COSMICv3.4.^14,44^ This analysis was performed using the *decompose_fit* function within SigProfilerAssignment v0.1.9.^45^ For each *de novo* signature, this function finds the optimal linear combination of COSMIC signatures that best reconstructs its mutational signature profile. A *de novo* signature could fail to decompose to any COSMIC signatures. We considered a *de novo* signature to be poorly decomposed if the reconstruction cosine similarity is below 0.90. In both cases, the *de novo* signature would be treated as potentially either novel or artifactual. The analysis was performed using the appropriate COSMIC reference set for the data type (WES or WGS) and reference genome (GRCh37 or GRCh38). Importantly, as previously done^31^, we excluded biologically irrelevant reference signatures (SBS7a, SBS7b, SBS7c, SBS7d, SBS10, and SBS12) before decomposing *de novo* signatures extracted from TCGA-BRCA and PCAWG Breast-AdenoCA.

### Assigning Signatures to Samples and Mutations

The number of samples in the input mutational catalog are assigned to *de novo* signatures using *cosmic_fit* in SigProfilerAssignment v0.1.9. We used *decompose_fit* to determine the activity of COSMICv3.4 reference signatures in the input samples.

### Manual inspection of putative mutations

TCGA-BRCA alignment BAM files were obtained from the GDC portal. We downloaded BAMs of PCAWG Breast-AdenoCA samples from the EU from the European Genome-Phenome Archive^46^ (https://ega-archive.org/datasets/EGAD00001002129). Duplicate reads were tagged by Picard MarkDuplicates using default parameters.^47^ We focused on the putative mutations exclusive to an individual variant caller that are in the same trinucleotide context as the caller-specific mutational signature, so SAMtools v1.13^24^ was used to subset BAMs. Reads were visualized on IGV (v2.16.1)^48^ either before MarkDuplicates (as the case for TCGA-BRCA) or after (for PCAWG Breast-AdenoCA). Reads were sorted by base, colored by read strand, and viewed as pairs.

### Statistical Analysis and Data Visualization

Cosine similarity was used as the primary metric to compare sample profiles and mutational signature profiles. Pearson’s correlation coefficient *r* was used as an alternative similarity metric. Variant allele frequency (VAF) was calculated as the number of reads supporting the alternate allele divided by the total number of reads at that base site. VAF distributions were visualized empirically as histograms using a bin size of 50. Spearman’s rank correlation was used to assess the relationship between sample profile similarity and number of SBS. All plots were generated using custom scripts in Python.

## Data and Code Availability

Public TCGA MC3 data are available from cBioPortal^49^ (https://www.cbioportal.org/), and harmonized GDC data can be accessed via the GDC Data Portal^50^ (https://portal.gdc.cancer.gov/). PCAWG VCFs are available through the ICGC Data Portal^42^ (https://dcc.icgc.org/releases/PCAWG) and BRCA-EU BAMs can be accessed through the European Genome-Phenome Archive^46^ (https://ega-archive.org/datasets/EGAD00001002129). Access to controlled data requires approval from the respective data access committees (e.g., eRA for TCGA, ICGC-DACO for PCAWG). The code used for all analyses and figure generation is available upon request.

## Acknowledgments/Funding

We thank Dr. Francisco Martínez-Jiménez (Vall d’Hebron Institute of Oncology, Barcelona, Spain) for generously sharing the SAGE WGS reanalysis for the ICGC samples from Martínez-Jiménez et al. (2023). M.D.-G. was awarded a fellowship within the “Generación D” initiative, Red.es, Ministerio para la Transformación Digital y de la Función Pública, for talent attraction (C005/24-ED CV1), funded by the European Union NextGenerationEU funds, through PRTR. M.D.-G. is hosted by the Centro Nacional de Investigaciones Oncológicas (CNIO), which is supported by the Instituto de Salud Carlos III and recognized as a “Severo Ochoa” Centre of Excellence (CEX2024-001442-S) by the Spanish Ministry of Science and Innovation (MCIN/AEI/10.13039/501100011033).

M.K. was supported by the Cancer Biology, Informatics, and Omics Training Grant (T32CA067754) and the Merkin Fellowship through the University of California San Diego. This work was supported by the US National Institute of Health grants R01ES032547, R01ES036931, R01CA269919, R01CA296974, P01CA281819, and U01CA290479 to L.B.A. as well as by L.B.A.’s Packard Fellowship for Science and Engineering and the UC San Diego Sanford Stem Cell Institute. The computational analyses reported in this manuscript have utilized the Triton Shared Computing Cluster at the San Diego Supercomputer Center of UC San Diego. The funders had no roles in study design, data collection and analysis, decision to publish, or preparation of the manuscript.

## SUPPLEMENTARY FIGURE LEGENDS

**Figure S1. Workflow for consensus calling and variant caller effects on WES mutational signature analysis.**

**(A)** Data processing workflow for comparison of consensus-called public TCGA WES data. Legacy and harmonized public TCGA datasets were filtered for PASS variants and intersected to identify 8,908 shared primary tumor samples across 32 cancer types. Mutational matrices were generated at both SBS-96 and SBS-192 resolutions using SigProfilerMatrixGenerator, followed by signature extraction using SigProfilerExtractor.

**(B)** Parallel curation workflow for assessing caller-specific artifacts in TCGA-BRCA. Controlled primary tumors (*n* = 736) shared between legacy and harmonized datasets were also used here. Mutations called by individual variant callers (MuTect2, VarScan2, MuSE) were processed at SBS-96 resolution.

**Figure S2. Workflow for variant caller effects on WGS mutational signature analysis.** The data curation strategy to assess the impact of variant calling methodology on signature identification and characterization across different mutational contexts in PCAWG whole genomes. Two sources of variants data were evaluated: (1) the original dataset (*n* = 2,583) comprised of consensus (2+/4) PASS mutations, where variants are called if detected by at least 2 of 4 variant callers (MuTect, MuSE, CaVEMan, DKFZ), along with PCAWG-filtered PASS mutations from each individual caller, termed "low support"^14^; and (2) a re-called dataset (*n* = 2,376) by the SAGE variant caller^26^ and filtered for PASS mutations. Samples were matched across all six variant calling approaches and restricted to 15 cancer types with >50 samples per cancer type, resulting in 1,857 samples. For each of the six variant calling approaches (PCAWG Consensus 2+/4, MuSE, CaVEMan, MuTect, DKFZ, and SAGE), mutational profiles were generated at four different resolutions (SBS-96, SBS-288, SBS-1536, and SBS-4608) using SigProfilerMatrixGenerator. *De novo* mutational signature extraction was then performed using SigProfilerExtractor.

**Figure S3. Poisson resampling of harmonized TCGA consensus SBS-96 mutational profiles.** Violin plots comparing 1,000 Poisson resampled replicates per sample to their original harmonized TCGA WES SBS-96 profiles (*n* = 8,908). The x-axis is plotted on a log_10_ scale, with labels representing raw mutation count of the original samples. The dark red dashed line marks the minimum mutation burden where 90% of simulations maintain a cosine similarity ≥0.95 with their original profiles. The bright red line indicates the threshold where 95% of simulations achieve this concordance.

**Figure S4. SBS-96 and SBS-192 *de novo* extraction selection plot in legacy and harmonized TCGA consensus datasets.** Selection plots from SigProfilerExtractor showing the optimal number of *de novo* mutational signatures across 32 TCGA cancer types. The plots demonstrate the effect of consensus-calling pipeline differences and extended mutational context (SBS-192) on WES signature extraction stability. Each subplot represents a different TCGA cancer type with sample sizes indicated (*n* = total samples; mutation counts in legacy and harmonized datasets). The x-axis shows the total number of signatures tested (range: 1 to 10), and the y-axis shows the average stability metric. Four different lines are plotted: Legacy SBS-96 signature solutions (blue diamond), Legacy SBS-192 signature solutions (blue circle), harmonized SBS-96 signature solutions (green diamond), and harmonized SBS-192 signature solutions (yellow circle). Squares indicate the suggested solution based on SigProfilerExtractor’s automated selection criteria, while stars indicate the most stable solution. Dotted horizontal lines indicate stability threshold of 0.80.

**Figure S5. Caller-specific SBS-96 mutational signatures in TCGA-BRCA are driven by variant caller’s artifactual mutations.**

**(A)** The mutational profile of a representative exome sample called by VarScan2 exhibits a prominent peak in the A[T>C]G context, mirroring a *de novo* signature extracted from the VarScan2 call set and from mutations unique to VarScan2. Integrative Genomics Viewer (IGV) visualization of a representative A[T>C]G site strongly suggests the VarScan2-exclusive call is a false positive.

**(B)** The profile of a representative exome sample called by MuSE resembles the MuSE-specific signature, particularly in the C[C>T]A context. IGV visualization suggests these mutations are technical artifacts.

**Figure S6. SBS-96 *de novo* extraction selection plot in PCAWG consensus variant calling and single variant callers.** Selection plots from SigProfilerExtractor comparing the optimal number of mutational signatures identified across different variant calling approaches for 15 cancer types in PCAWG WGS. Each subplot represents a different cancer type, with the x-axis showing the total number of signatures tested (range: 1 to 25) and the y-axis showing the average stability metric. Colored lines represent different variant calling approaches: PCAWG consensus (2+/4; grey), MuSE (blue), CaVEMan (orange), SAGE (green), MuTect (red), and DKFZ (purple). The grey shaded rectangle indicates the suggested solution determined by SigProfilerExtractor for the consensus variant calling approach. The plots demonstrate how variant calling methodology affects signature extraction stability and the optimal number of signatures identified. Differences in stability patterns and suggested solutions across variant callers highlight the impact of technical variation on mutational signature analysis. Cancer type abbreviations are shown at the top of each subplot.

**Figure S7. Caller-specific SBS-96 mutational signatures in PCAWG Breast-AdenoCA are driven by variant caller’s artifactual mutations.**

**(A)** The mutational profile of a representative whole-genome sequenced sample called by MuTect exhibits a subtle excess in G[C>T]T compared to consensus. This pattern across many samples manifests as a MuTect caller-specific signature that is reproduced from MuTect-exclusive calls. IGV visualization of a representative G[C>T]T site confirms that false positives drive this peak.

**(B)** The profile of a representative whole-genome sequenced MuTect sample is strongly enriched in the G[T>G]G context, resembling another artifactual signature extracted from the MuTect call set and mutations exclusive to MuTect. IGV visualization suggests these G[T>G]G mutations are technical artifacts.

**(C)** A representative whole-genome sequenced sample called by SAGE displays a C[C>G]T peak that exceeds the consensus, matching the SAGE caller-specific signature and the signature derived from SAGE-exclusive mutations. Visualization of a representative C[C>G]T site confirms these calls are technical artifacts.

**Figure S8. Consensus variant calling produces consistent *de novo* mutational signatures in SBS-96.** Heatmaps displaying the cosine similarity between the PCAWG Breast-AdenoCA SigProfilerExtractor suggested solution derived from the intersection of all five variant callers and stable solutions obtained from individual callers or different consensus methods. Results for the commonly used SBS-96 mutational context, demonstrating that our findings hold across different mutational resolutions. Each extraction’s suggested number of signatures and the total mutation count are listed below the corresponding tick labels.

**Figure S9. Consensus 2+/5 *de novo* signature extraction across different tools in SBS-96 WGS Breast-AdenoCA.** Comparison of *de novo* mutational signatures extracted from PCAWG Breast-AdenoCA samples using consensus variant calling (2+/5 threshold) across three signature extraction tools: SigProfilerExtractor (SPE; left column), SignatureToolsLib (STL; middle column), and MuSiCal (right column). Each row represents signatures identified by the different tools, with signature names and cosine similarities to the SPE signature shown above each profile. Bar plots display the percentage of mutations (y-axis) across the 96 trinucleotide contexts (x-axis), color-coded by mutation type: C>A (cyan), C>G (black), C>T (red), T>A (grey), T>C (green), and T>G (pink). "Not detected" indicates a signature is extracted by other tools but missed by a particular extraction tool.

**Figure S10. SignatureToolsLib *de novo* SBS-96 extractions across variant calling methods forced at the same *k*.** *De novo* signatures extracted using SignatureToolsLib (STL) across multiple variant calling strategies: Intersect 5 (variants detected by all five callers), Consensus 2+/5, Consensus 2+/3, pairwise caller combinations (MuTect + SAGE, MuTect + MuSE), and five individual variant callers (MuTect, MuSE, SAGE, CaVEMan, DKFZ). All extractions were forced to use the number of signatures suggested by “Intersect 5” to enable direct comparison across calling methods. Selection plots (top) show stability metrics, with the forced solution indicated with a grey vertical bar. Signature profiles display the percentage of mutations across 96 trinucleotide contexts, color-coded by mutation type. Gray rectangles indicate signatures identified by other variant calling approaches but missed by a particular variant calling method (cosine similarity <0.85). Artifactual signatures showed high concordance (>0.95) between STL and SigProfilerExtractor (SPE), including STL SAGE S6 with SPE SAGE SBS96H (cosine 0.9960) and STL MuTect S1 with SPE MuTect SBS96J (cosine 0.9955), demonstrating consistent identification of technical artifacts across extraction tools.

**Figure S11. MuSiCal *de novo* SBS-96 extractions across variant calling methods forced at the same *k*.** *De novo* signatures extracted using MuSiCal across multiple variant calling strategies: Intersect 5 (variants detected by all five callers), Consensus 2+/5, Consensus 2+/3, pairwise caller combinations (MuTect + SAGE, MuTect + MuSE), and five individual variant callers (MuTect, MuSE, SAGE, CaVEMan, DKFZ). All extractions were forced to use the same number of signatures suggested by “Intersect 5” to enable direct comparison across calling methods. Selection plots (top) show stability metrics. MuSiCal automatically picks an optimal number of signatures, marked by a grey vertical bar. The forced number of signatures is highlighted with a purple vertical bar. Signature profiles display the percentage of mutations across 96 trinucleotide contexts, color-coded by mutation type. Gray rectangles indicate signatures identified by other variant calling approaches but absent in a particular calling method (cosine similarity <0.85). Artifactual signatures showed high concordance (>0.95) between MuSiCal and SigProfilerExtractor (SPE), including MuSiCal SAGE SBS96F with SPE SAGE SBS96H (cosine 0.9890) and MuSiCal MuTect SBS96D with SPE MuTect SBS96J (cosine 0.9810), demonstrating consistent identification of technical artifacts across extraction tools.

**Figure S12. Legacy and harmonized TCGA consensus datasets’ SBS-192 mutational profile and *de novo* signature cosine similarities.**

**(A)** Histogram showing the number of TCGA primary tumors at each bin for SBS-192 sample profile cosine similarity between legacy and harmonized consensus calling datasets.

**(B)** Scatter plot comparing a sample’s SBS mutation count in the harmonized dataset (x-axis; log-scale) and the cosine similarity (y-axis) between legacy and harmonized releases. Each dot corresponds to an individual tumor. Samples with cosine similarity below 0.85 are highlighted in orange. An accompanying box plot on the right illustrates the distribution of cosine similarities.

**(C)** *Top*: Stacked bar plot showing the proportion of samples per cancer type within three tiers of cosine similarity (≥0.95, [0.85,0.95), and <0.85). Cancer types are sorted by the proportion of samples in the ≥0.95 bin. *Bottom*: Snake plot showing the mutational burden (SBSs per 55 Megabases (Mb); assuming that the exome is 55 MB) for each sample in the harmonized dataset, grouped by cancer type. Horizontal lines indicate the median mutational burden for the harmonized (black) and legacy (red) datasets.

**(D)** Stacked bar plot showing the number of nearly identical, similar, and dissimilar *de novo* extracted SBS-192 signatures between legacy and harmonized datasets for each cancer type, at a forced number of signatures, which is often the suggested *k* in that cancer type. The default normalization method of Gaussian mixture model was used. Dissimilar signatures that were altered by the harmonization process are highlighted in orange. i.e., Neck Squamous Cell Carcinoma (HNSC) and Acute Myeloid Leukemia (LAML).

**Figure S13. Comparison of individual versus PCAWG consensus variant calling for extended SBS-288 context mutational signature extractions.**

**(A)** Scatter plots showing the cosine similarity of SBS-288 sample profile between PCAWG consensus (2+/4) and five individual variant callers, in relation to SBS mutation count (x-axis; log-scale). Each dot corresponds to an individual tumor. Samples with cosine similarity below 0.85 are highlighted in orange.

**(B)** *Top*: Stacked bar plot showing the proportion of samples per cancer type per variant caller within three tiers of cosine similarity (≥0.95, [0.85,0.95), and <0.85). Cancer types are sorted by the proportion of samples in the ≥0.95 bin averaged across the five variant callers. *Bottom*: Snake plot showing the mutational burden (SBSs per 2,800 Megabases (Mb); assuming the genome is 2,800 MB) of each sample determined by each individual variant caller. Black horizontal lines indicate the median mutational burden for the datasets.

**(C)** Stacked bar plot showing the number of nearly identical, similar, and dissimilar *de novo* extracted SBS-288 signatures between consensus (2+/4) and individual variant callers (see online methods for solution picking in WGS extractions). The number of caller-specific signatures in a cancer type is highlighted in lighter hues.

**Figure S14. Comparison of individual versus PCAWG consensus variant calling for extended SBS-1536 context mutational signature extractions.**

**(A)** Scatter plots showing the cosine similarity of SBS-1536 sample profile between PCAWG consensus (2+/4) and five individual variant callers, in relation to SBS mutation count (x-axis; log-scale). Each dot corresponds to an individual tumor. Samples with cosine similarity below 0.85 are highlighted in orange.

**(B)** *Top*: Stacked bar plot showing the proportion of samples per cancer type per variant caller within three tiers of cosine similarity (≥0.95, [0.85,0.95), and <0.85). Cancer types are sorted by the proportion of samples in the ≥0.95 bin averaged across the five variant callers. *Bottom*: Snake plot showing the mutational burden (SBSs per 2,800 Megabases (Mb); assuming the genome is 2,800 MB) of each sample determined by each individual variant caller. Black horizontal lines indicate the median mutational burden for the datasets.

**(C)** Stacked bar plot showing the number of nearly identical, similar, and dissimilar *de novo* extracted SBS-1536 signatures between consensus (2+/4) and individual variant callers (see online methods for solution picking in WGS extractions). The number of caller-specific signatures in a cancer type is highlighted in lighter hues.

**Figure S15. Comparison of individual versus PCAWG consensus variant calling for extended SBS-4608 context mutational signature extractions.**

**(A)** Scatter plots showing the cosine similarity of SBS-4608 sample profile between PCAWG consensus (2+/4) and five individual variant callers, in relation to SBS mutation count (x-axis; log-scale). Each dot corresponds to an individual tumor. Samples with cosine similarity below 0.85 are highlighted in orange.

**(B)** *Top*: Stacked bar plot showing the proportion of samples per cancer type per variant caller within three tiers of cosine similarity (≥0.95, [0.85,0.95), and <0.85). Cancer types are sorted by the proportion of samples in the ≥0.95 bin averaged across the five variant callers. *Bottom*: Snake plot showing the mutational burden (SBSs per 2,800 Megabases (Mb); assuming the genome is 2,800 MB) of each sample determined by each individual variant caller. Black horizontal lines indicate the median mutational burden for the datasets.

**(C)** Stacked bar plot showing the number of nearly identical, similar, and dissimilar *de novo* extracted SBS-4608 signatures between consensus (2+/4) and individual variant callers (see online methods for solution picking in WGS extractions). The number of caller-specific signatures in a cancer type is highlighted in lighter hues.

**Figure S16. Consensus variant calling produces consistent *de novo* mutational signatures in SBS-288.** Heatmaps displaying the cosine similarity between the PCAWG Breast-AdenoCA SigProfilerExtractor suggested solution derived from the intersection of all five variant callers and stable solutions obtained from individual callers or different consensus methods. Results for the extended SBS-288 context, demonstrating that our findings hold across different mutational resolutions. Each extraction’s suggested number of signatures and the total mutation count are listed below the corresponding tick labels.

