## Supplementary Figures 1-16 for "The Impact of Variant Calling on Substitution Mutational Signature Inference"

#### Fig. S1 Workflow for consensus calling and variant caller effects on WES mutational signature analysis

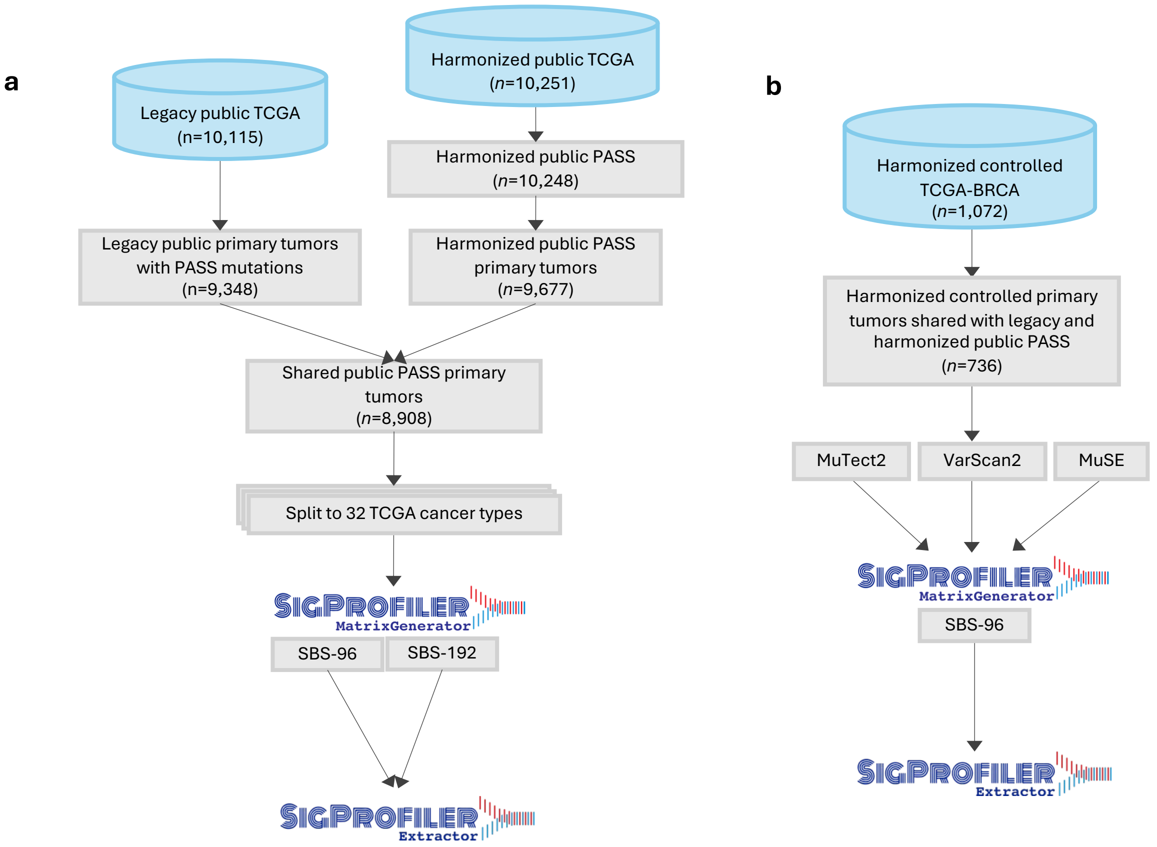

**Fig. S2 Workflow for variant caller effects on WGS mutational signature analysis**

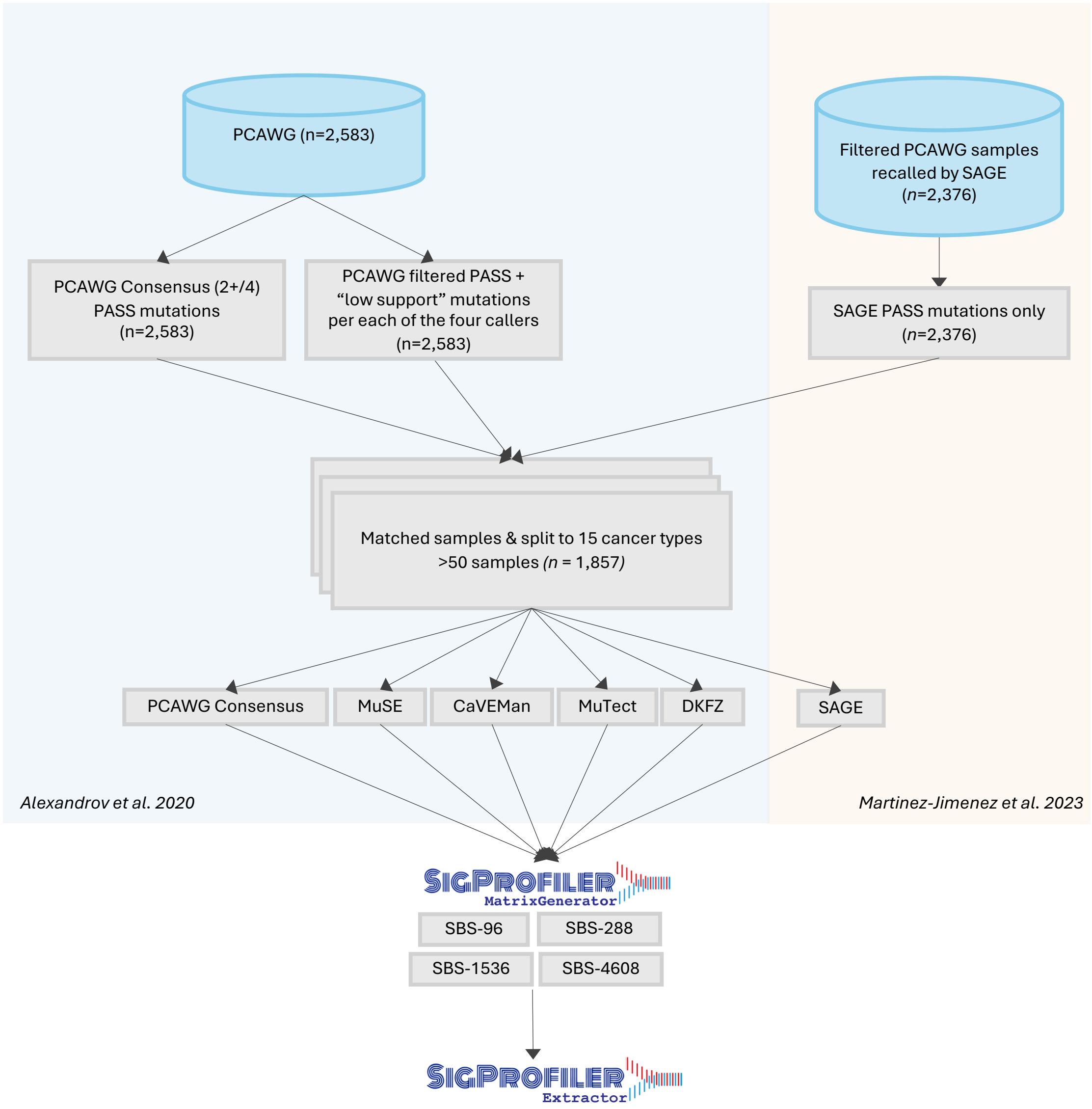

### Fig. S3 Poisson resampling of harmonized TCGA consensus SBS-96 mutational profiles.

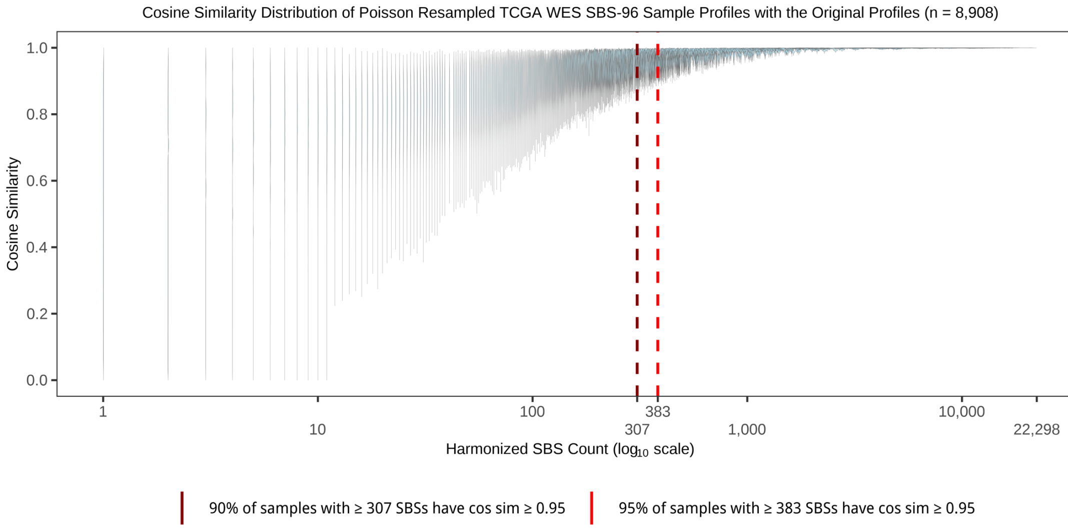

**Fig. S4 SBS-96 and SBS-192 *de novo* extraction selection plot in legacy and harmonized TCGA consensus datasets.**

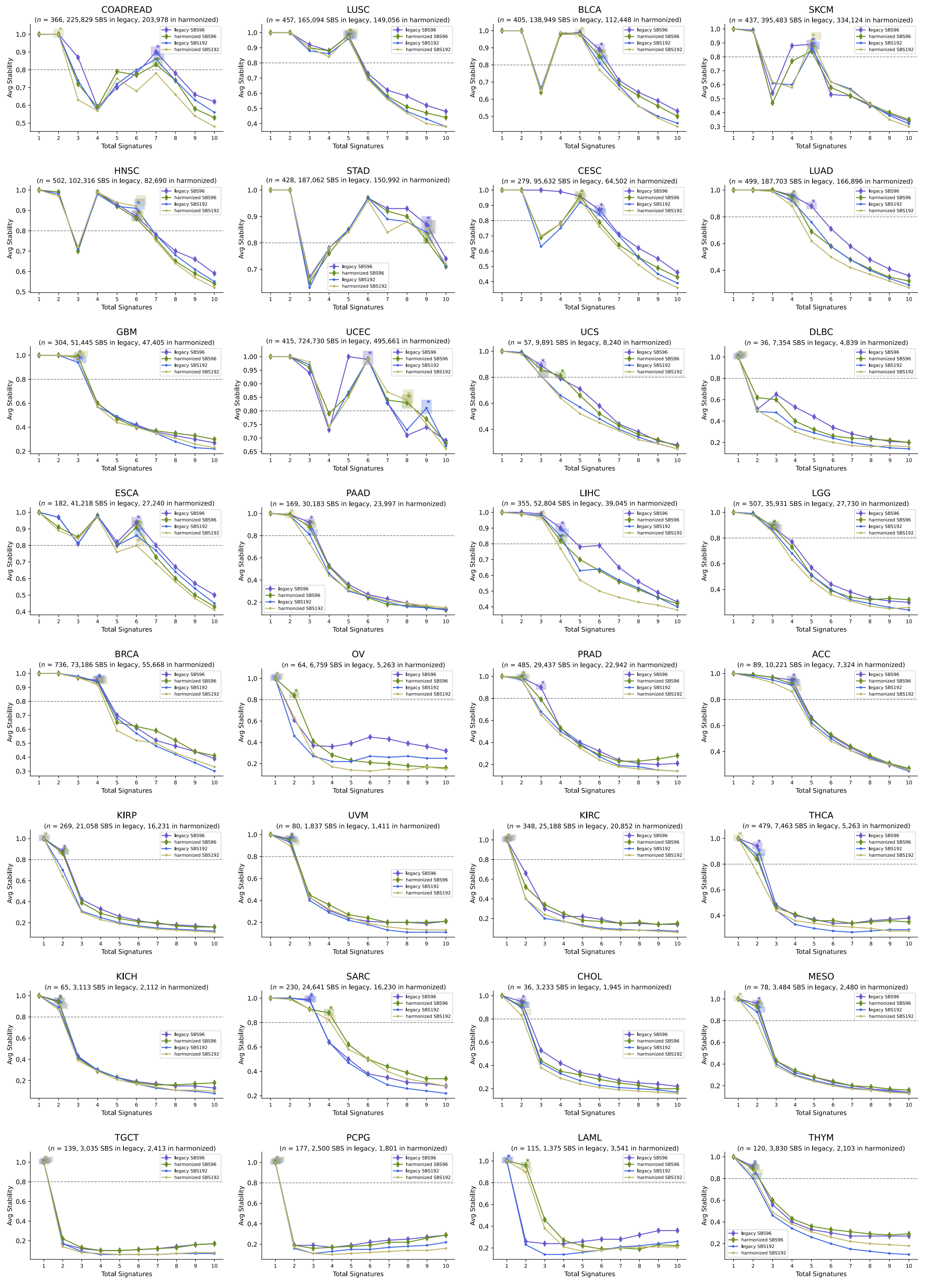

### Fig. S5 Caller-specific SBS-96 mutational signatures in TCGA-BRCA are driven by variant caller's artifactual mutations.

A

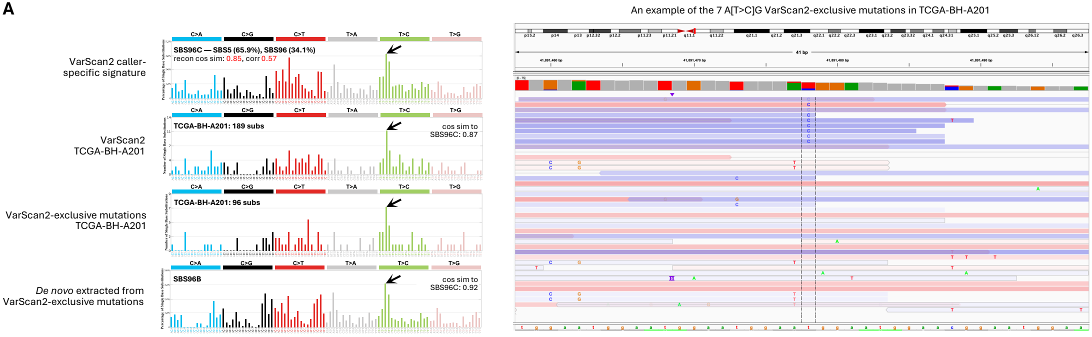


B

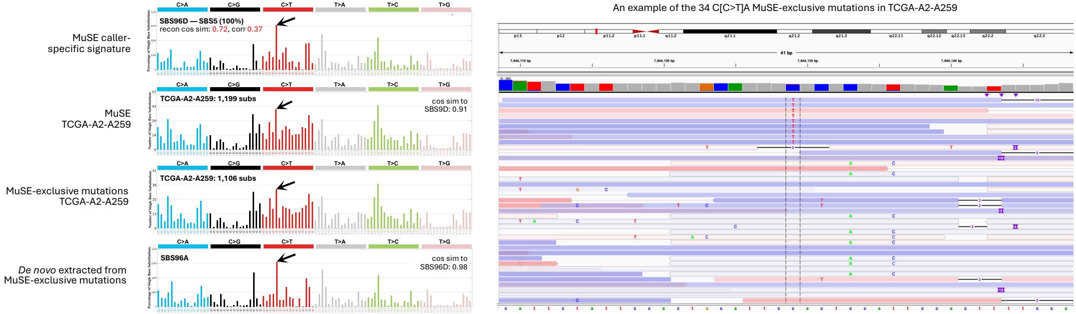

### Fig. S6 SBS-96 *de novo* extraction selection plot in PCAWG consensus variant calling and single variant callers.

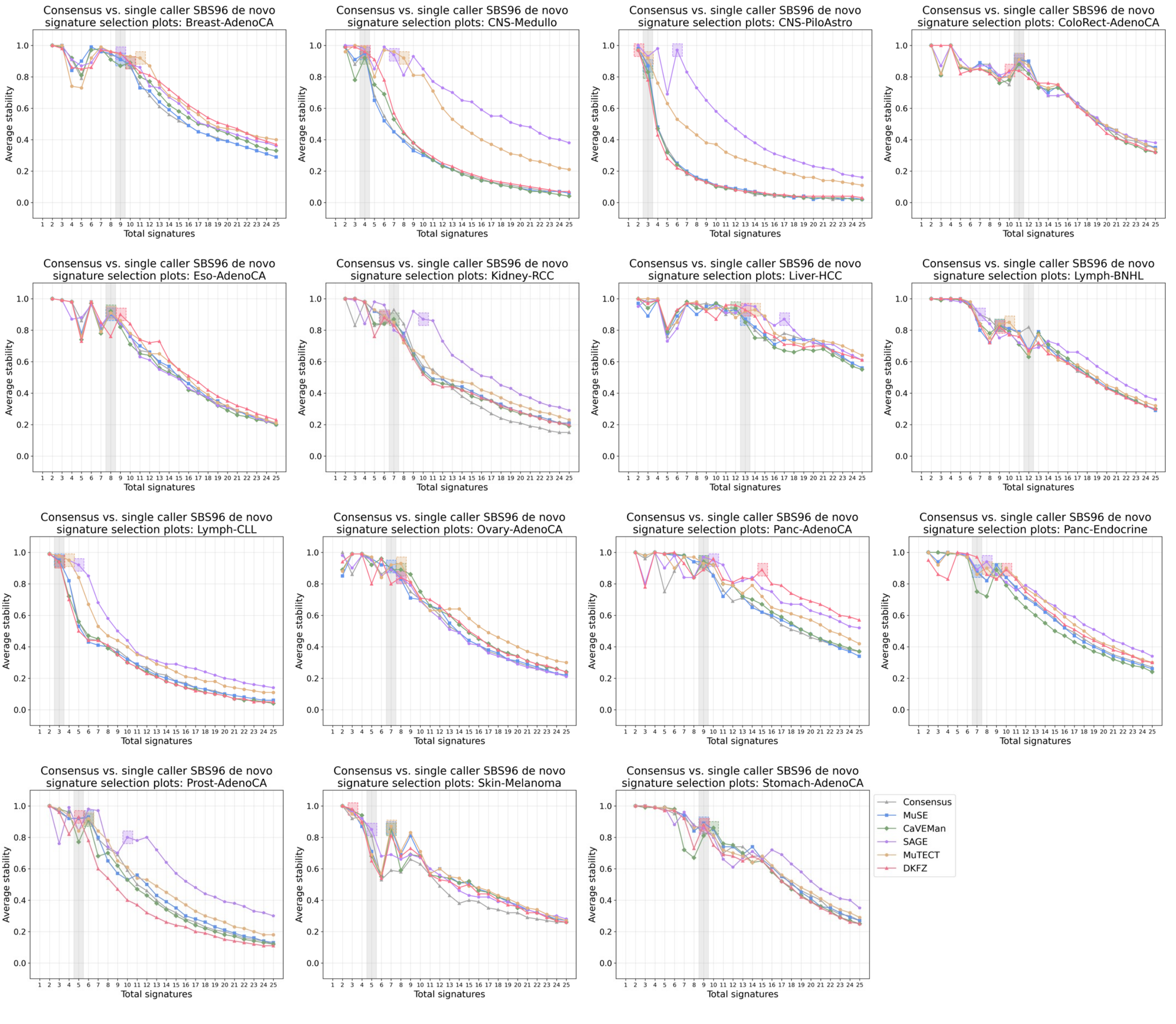

**Fig. S7 Caller-specific SBS-96 mutational signatures in PCAWG Breast-AdenoCA are driven by variant caller's artifactual mutations.**

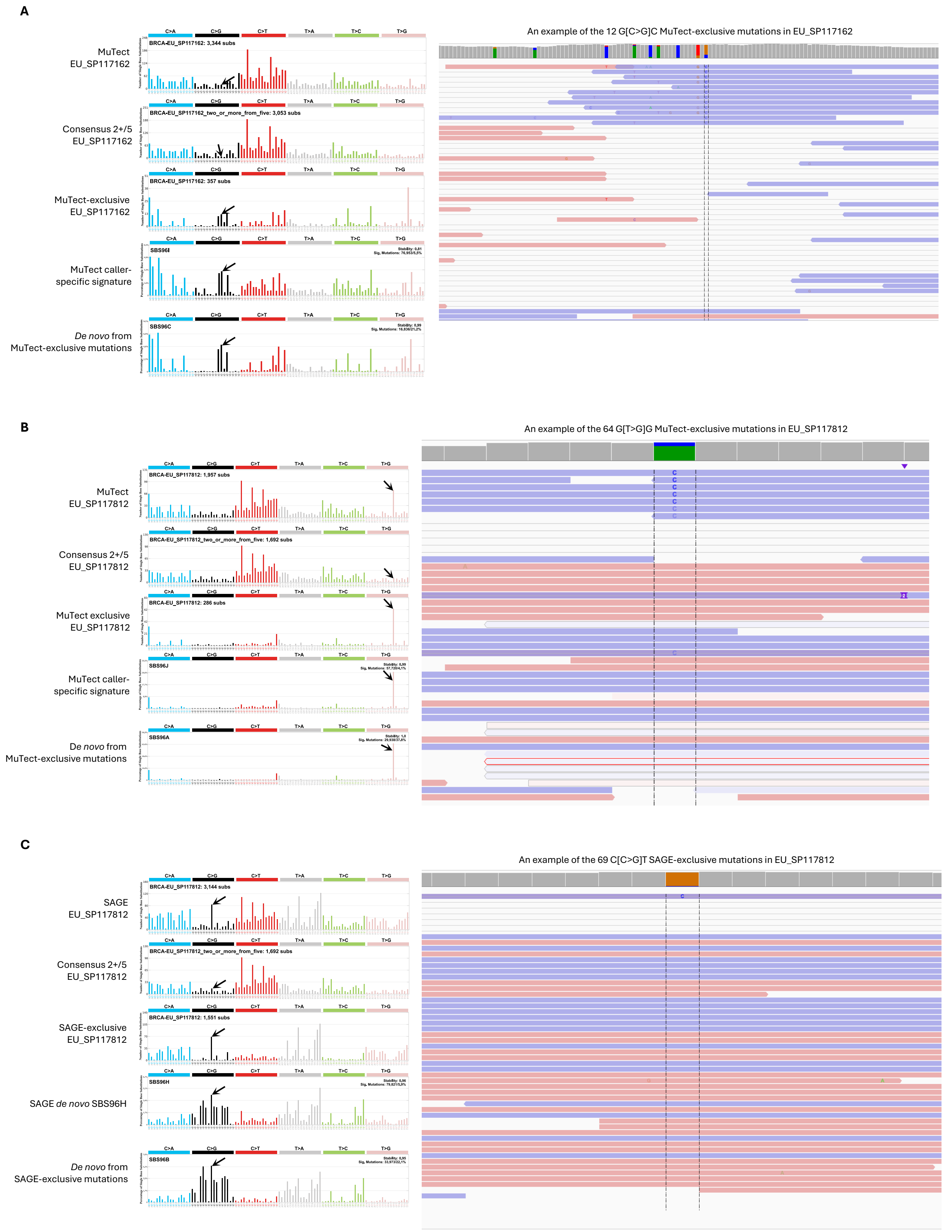

### Fig. S8 Consensus variant calling produces consistent *de novo* mutational signatures in SBS-96

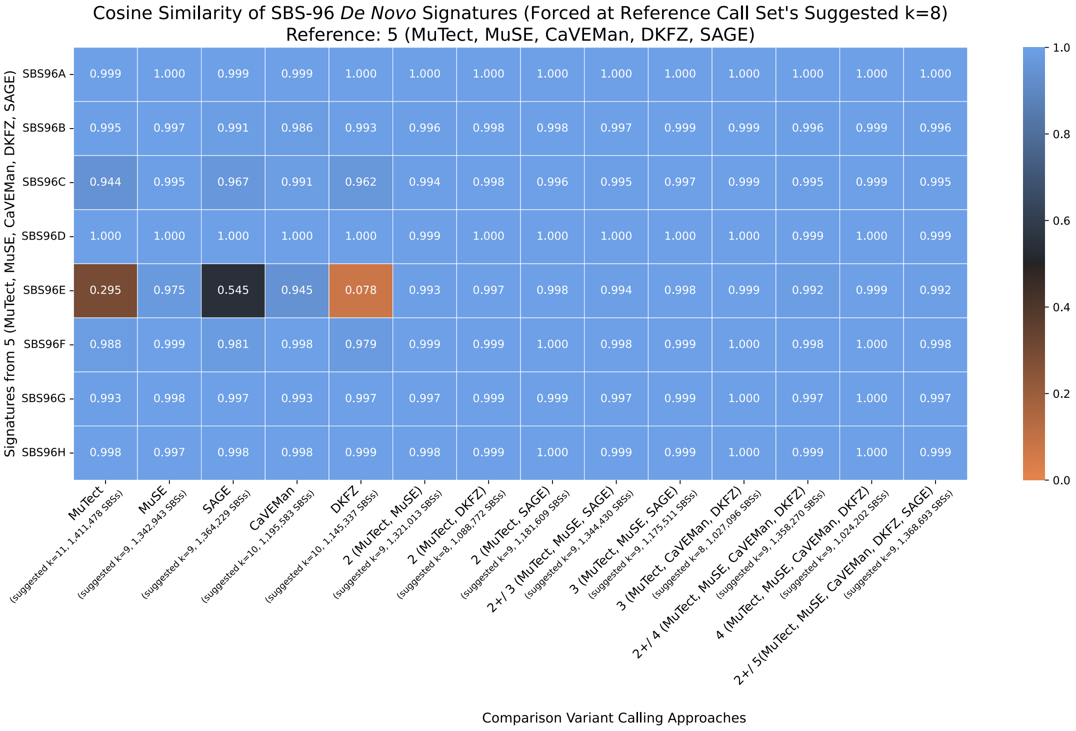

### Fig. S9 Consensus 2+/5 *de novo* signature extraction across different tools in SBS-96 WGS Breast-AdenoCA.

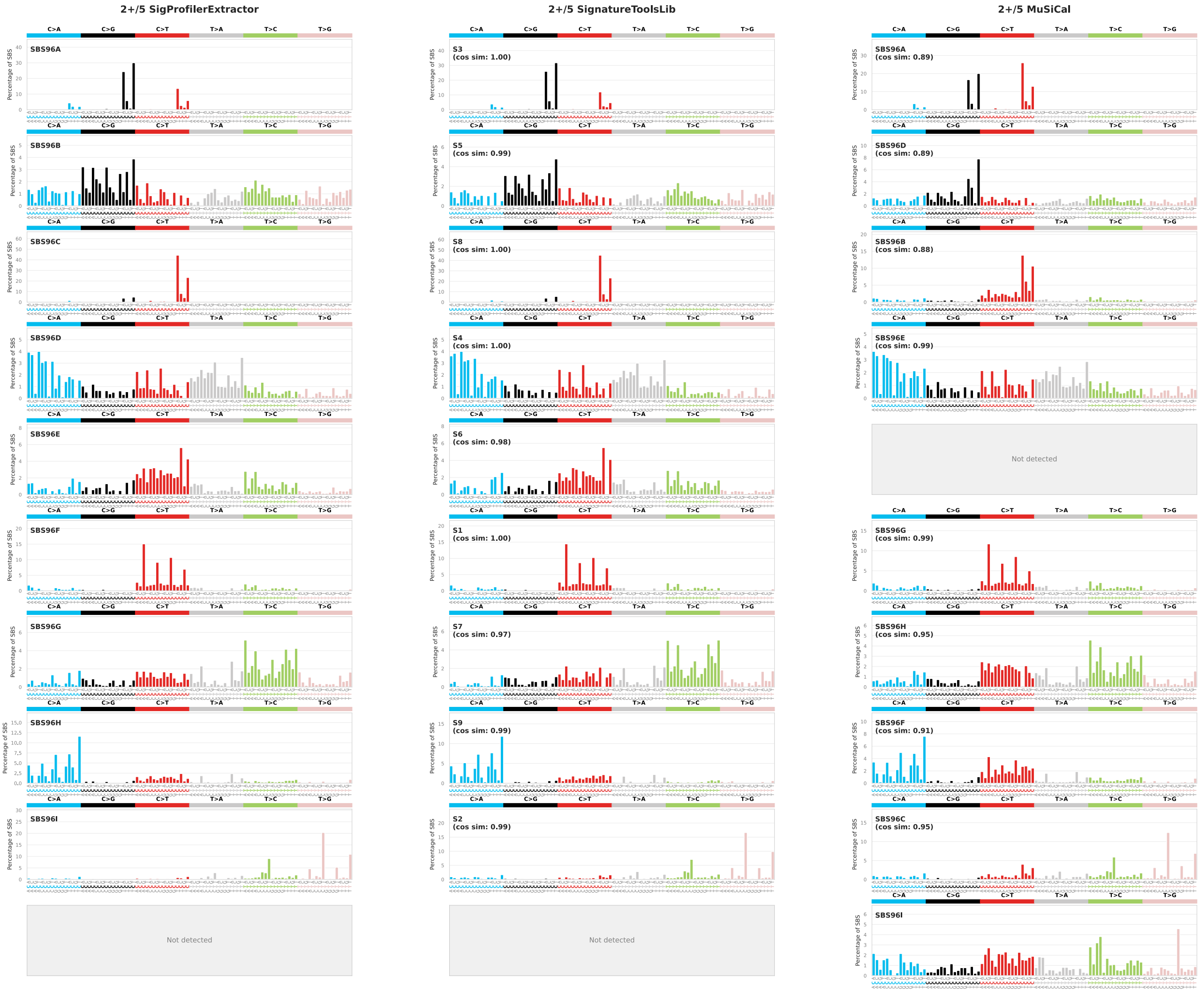

**Fig. S10 SignatureToolsLib *de novo* SBS-96 extractions across variant calling methods forced at the same *k*.**

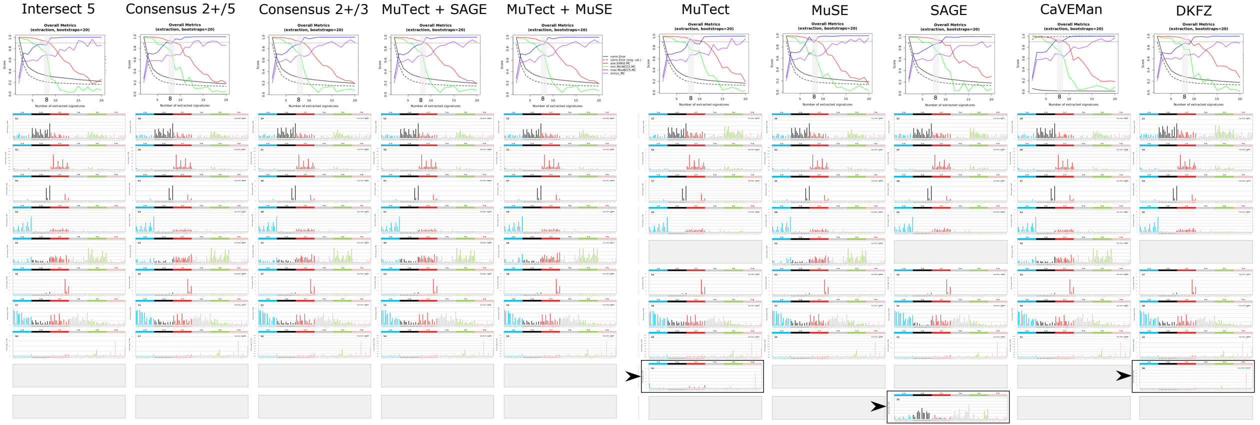

Fig. S11 MuSiCal *de novo* SBS-96 extractions across variant calling methods forced at the same *k*.

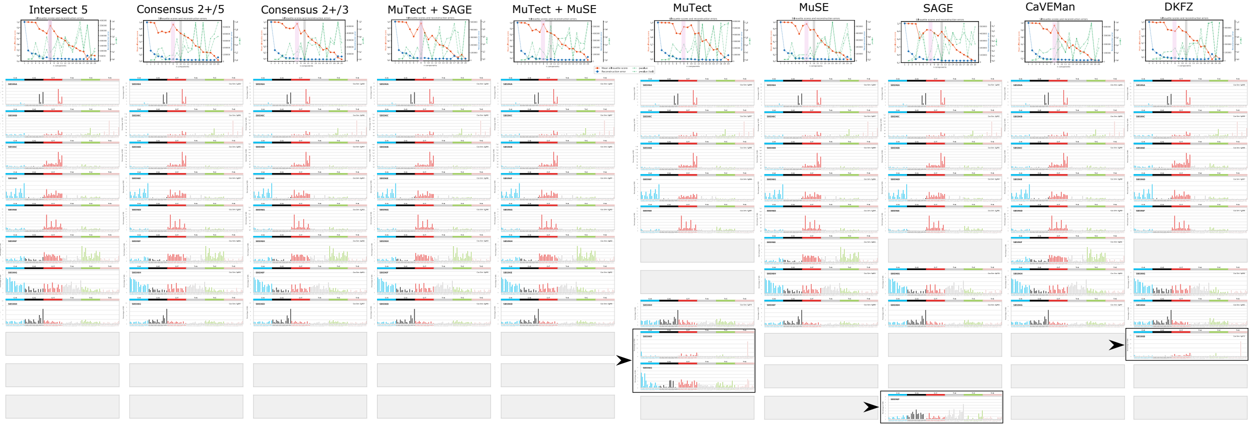

**Fig. S12 Legacy and harmonized TCGA consensus datasets' SBS-192 mutational profile and *de novo* signature cosine similarities.**

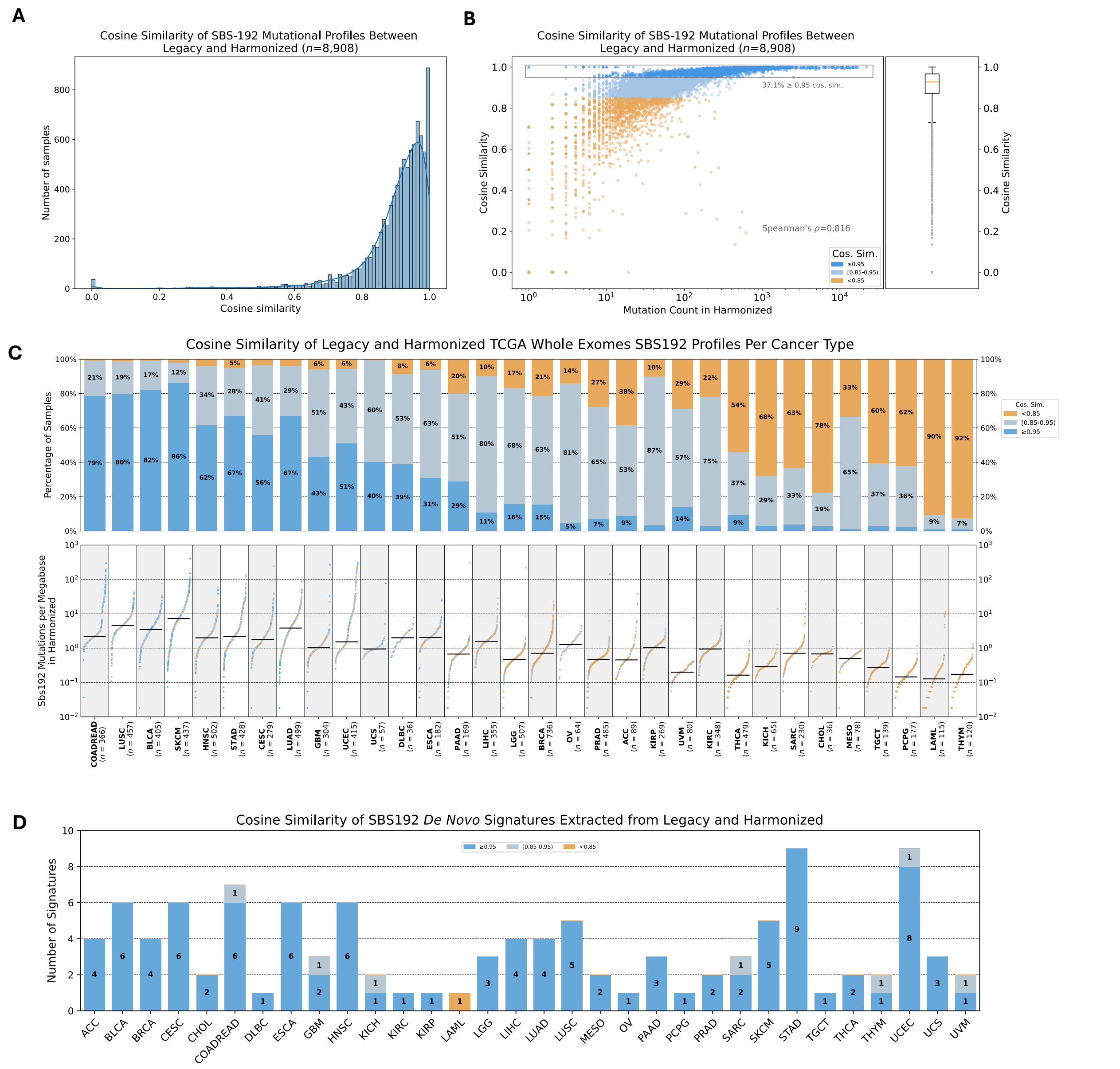

### Fig. S13 Comparison of individual versus PCAWG consensus variant calling for extended SBS-288 context mutational signature extractions

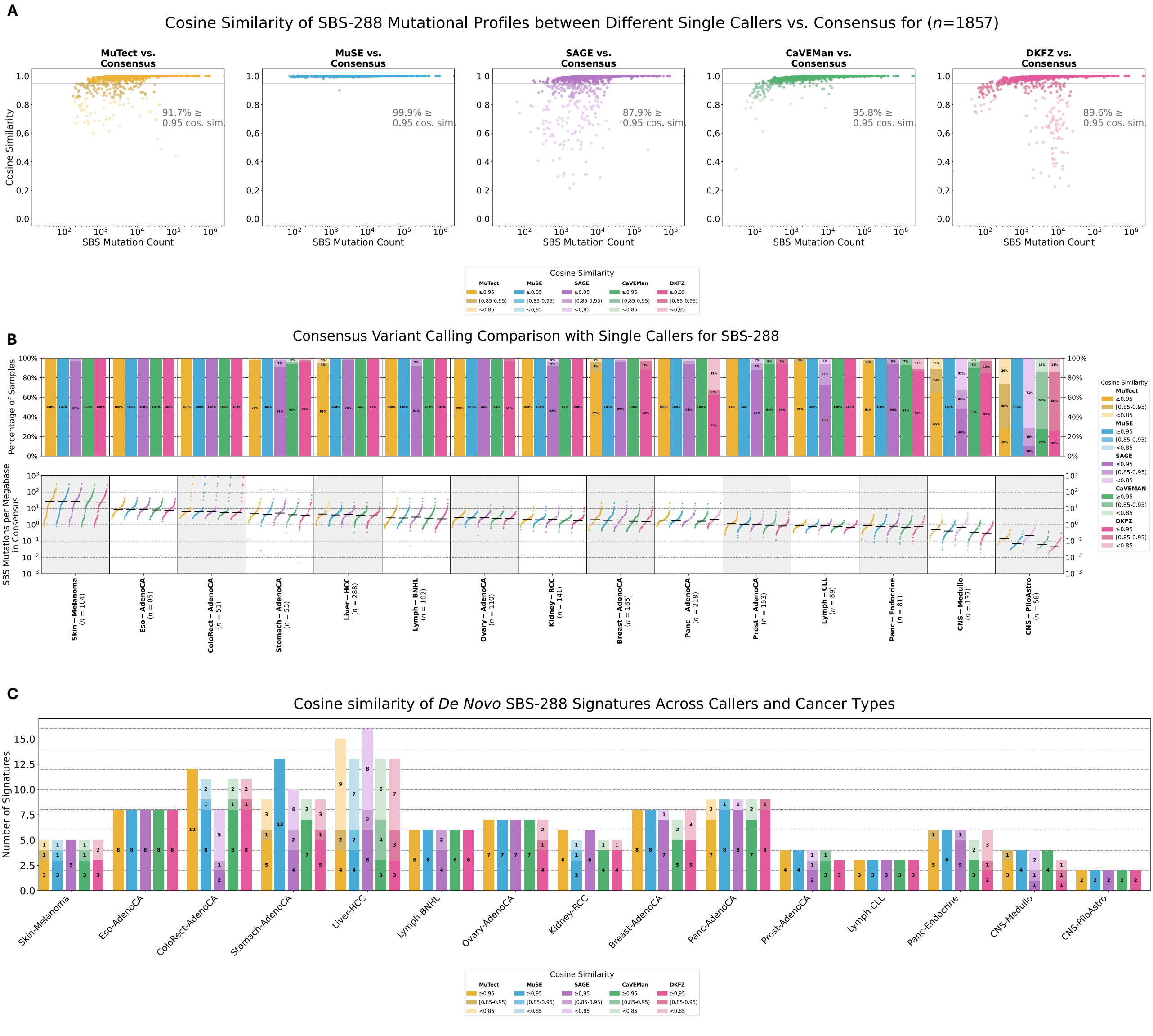

### Fig. S14 Comparison of individual versus PCAWG consensus variant calling for extended SBS-1536 context mutational signature extractions

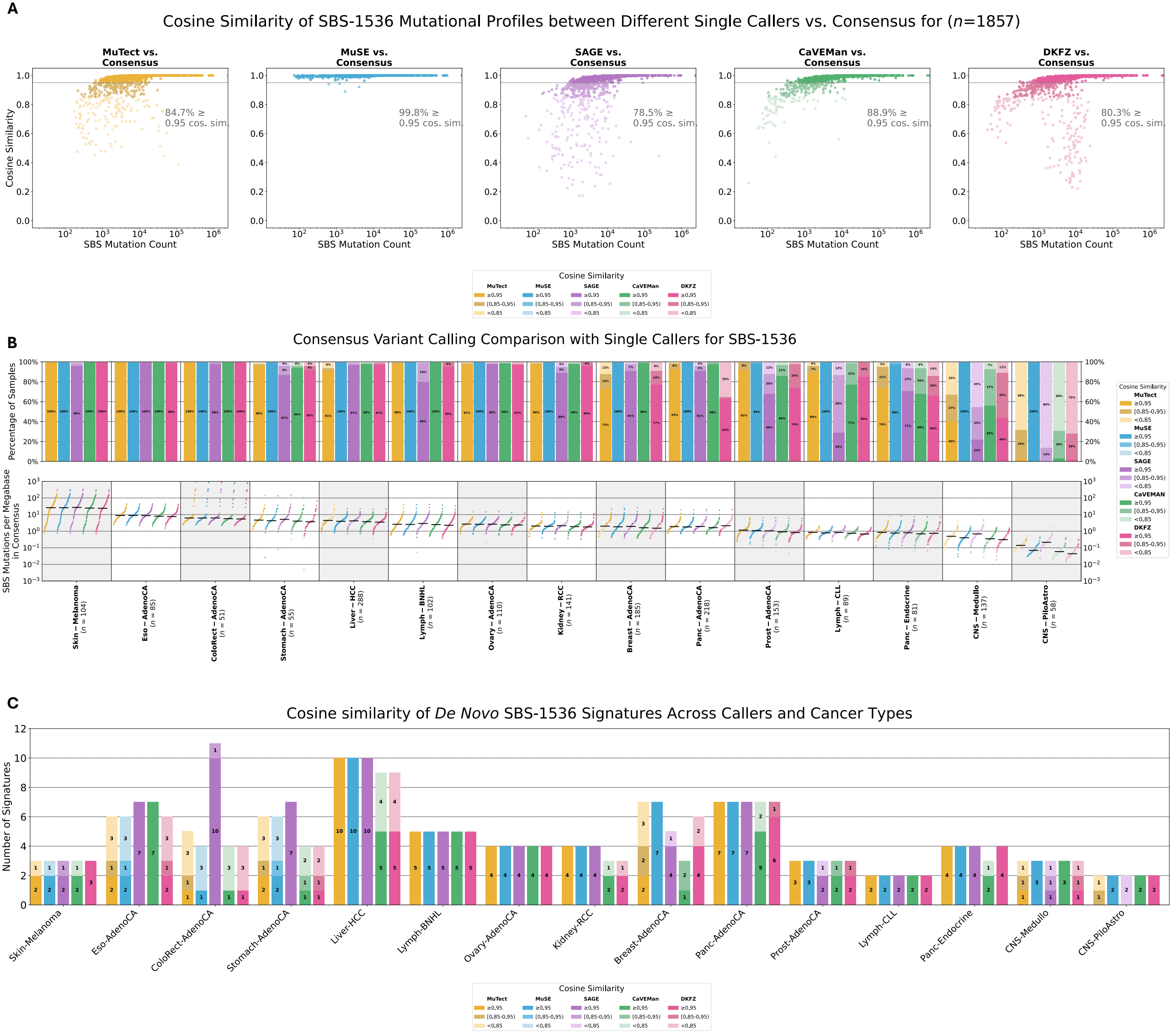

### Fig. S15 Comparison of individual versus PCAWG consensus variant calling for extended SBS-4608 context mutational signature extractions

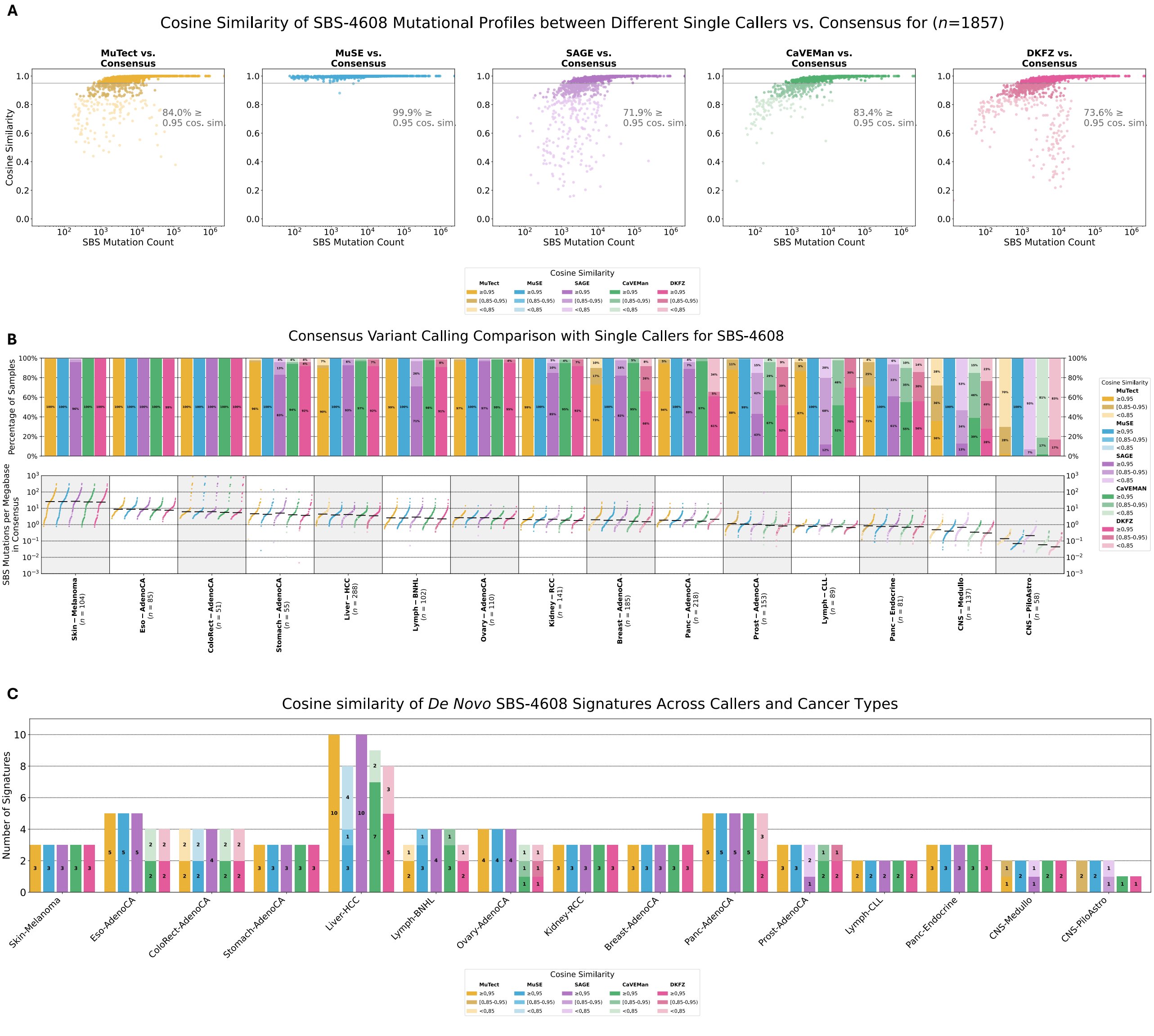

#### Fig. S16 Consensus variant calling produces consistent *de novo* mutational signatures in SBS-288

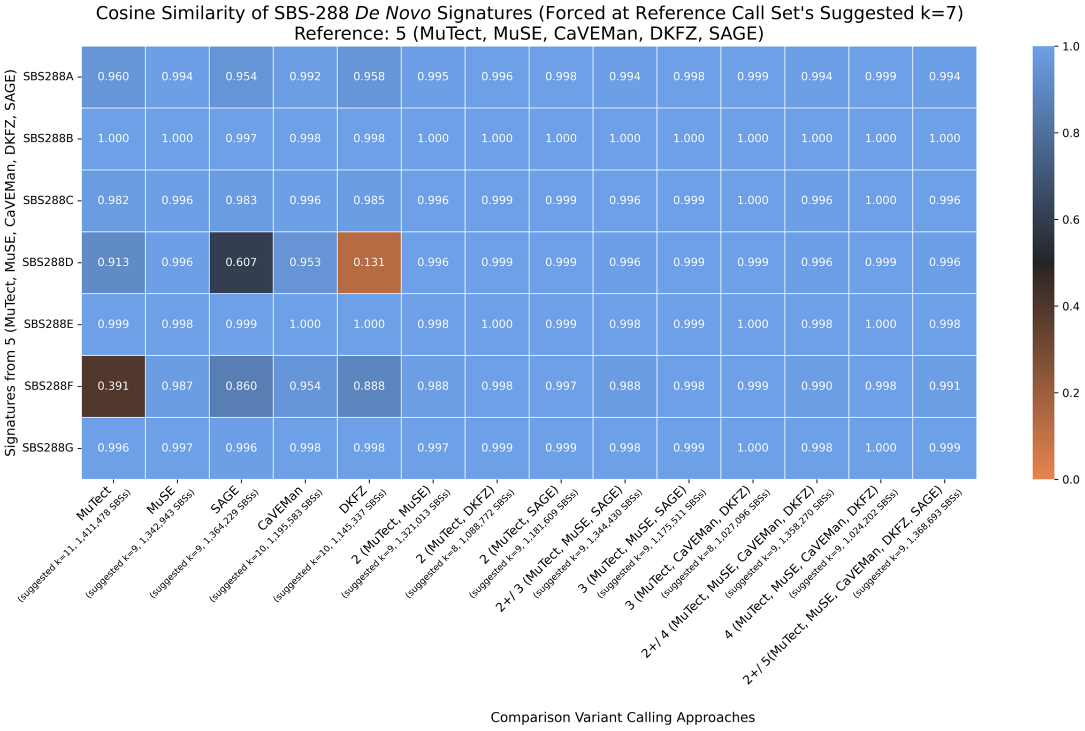
